# The Genomic Basis of Social Parasitism: A Geographical Mosaic of Behavioural, Chemical, and Environmental Adaptations in a Widespread Host–Parasite System

**DOI:** 10.1101/2025.05.24.655926

**Authors:** Maide Nesibe Macit, Erwann Collin, Maria Esther Nieto-Blazquez, Marion Kever, Maria Litto, Esther Jaitner, Markus Pfenninger, Barbara Feldmeyer, Susanne Foitzik

**Author notes:** shared first authors (Maide Nesibe Macit, Erwann Collin). shared last authors.

## Abstract

Coevolutionary dynamics in host–parasite systems are driven by reciprocal selection and environmental pressures. When parasite and host are closely related and have similar evolutionary potentials, evolution may follow parallel trajectories, affecting the same traits and underlying genes. We investigated coevolution and its genomic basis in the dulotic ant parasite *Temnothorax americanus* and its host *T. longispinosus* across a broad climatic gradient using population genomics, genome-wide association and transcriptome analyses. Population genomics revealed a striking contrast: panmictic host populations versus structured parasite populations, consistent with geographic mosaic dynamics. Genomic responses to parasite prevalence were strongly asymmetric: hosts showed strong selection on immune and structural defence genes, potentially with pleiotropic social functions. Parasites exhibited weaker signals, often in regulatory genes linked to behavioural shifts critical for raiding. Both species displayed shared genomic signatures of climate adaptation (e.g., desiccation resistance, stress response), suggesting convergent physiological responses. Genes associated with host–parasite encounters (mechanosensation, circadian rhythms, venom) also showed parallel selection. Behavioural traits such as aggression showed limited genomic signals but potentially higher transcriptional plasticity. Associations with chemical traits revealed shared selection on genes involved in cuticular hydrocarbon biosynthesis and chemosensory perception, indicating evolutionary coupling of signal production and perception. Constitutive gene expression patterns diverged: host expression correlated with parasite prevalence, while parasite expression was more strongly linked to climate, reflecting contrasting regulatory pressures. Our study demonstrates how differing population structures, asymmetric reciprocal selection, and environmental context shape divergent genomic trajectories of coadaptation, reflecting distinct evolutionary architectures across a heterogeneous landscape.

## Introduction

Antagonistic interactions between hosts and parasites can lead to dynamic coevolutionary arms races where both parties continuously adapt and diversify in response to one another (Dawkins & Krebs, 1979; Hamilton, 1982; Nash et al., 2008; Tellier et al., 2014). These arms races occur at the population rather than species level, with the independence of local interactions shaped by the extent of gene flow among populations. According to the geographic mosaic of coevolution (Thompson, 1999, 2005), resulting arms races escalate at different rates across each mosaic piece, with variations in environmental conditions and community compositions leading to divergent local adaptations. Some traits may evolve in response to both antagonistic partners and abiotic conditions, where trade-offs may arise when both demands cannot be fulfilled simultaneously (Wolinska & King, 2009; Amandine et al., 2022). This presents two interrelated challenges: i) determining which observed changes in phenotypic traits result from the coevolutionary arms race versus environmental conditions, and ii) inferring the extent to which these processes are interdependent or linked through pleiotropic interactions (Thompson, 1999, 2005). Such interactions can promote or constrain trait expression in antagonistic relationships, occasionally resulting in counterintuitive outcomes (Clarke & Fraser, 2004; Gregory, 2009; Wisz et al., 2013; Oppold et al., 2016; Mahmud et al., 2017). Species population structure, shaped by historical events and ongoing gene flow, influences evolutionary cohesion and the available adaptive genetic variations (Whitlock, 2004; Durrett & Schweinsberg, 2005; Greischar & Koskella, 2007; Gupta & Vadde, 2019). Consequently, it also shapes the adaptive evolutionary trajectories of these species in response to environmental and biotic selection pressures across their geographic ranges (Aaltonen, 2002; Tigano & Friesen, 2016). To determine the genomic basis of these interacting evolutionary forces, a comprehensive analysis must consider them simultaneously.

Brood and social parasitism is a lifestyle characterised by behavioural manipulation and exploitation of host social traits (Hughes et al., 2012; Jamie & Kilner, 2017; Jongepier et al., 2015; Rojas-Ripari et al., 2021). Social parasites evade the costs associated with labour tasks such as brood care, nest construction, and foraging by exploiting their host’s workforce (Lenoir et al., 2001; Buschinger, 2009). Hosts can counter-evolve a range of defences to mitigate the fitness costs of parasitism, including avoidance behaviour, aggressive defences, and tolerance strategies limiting parasite-inflicted damages (Råberg et al., 2009; Feeney et al., 2014; Gibson & Amoroso, 2022). Recognition is a prerequisite for successful counteradaptation to detect and discriminate parasites from conspecifics or benign species (Langmore et al., 2011). Hosts of avian brood parasites may evade parasitism by identifying and immediately rejecting foreign eggs (Soler, 2017; Manna et al., 2019). Similarly, hosts of social insect parasites initiate defence responses upon identifying parasite-specific chemical cues (Delattre et al., 2012). Such cue recognition is facilitated through odorant receptors located in the antennal sensilla of insects (Hölldobler & Wilson, 1990; Ozaki et al., 2005). Odorant receptor genes have undergone expansion during the evolution of insect sociality, along with the emergence of complex chemical communication and the regulation of social organisation (McKenzie & Kronauer, 2018; Gautam et al., 2024). Social parasites have convergently lost many odorant receptor genes, particularly those linked to worker behaviour (Caminer et al., 2023; Harrison et al., submitted) due to relaxed selection on traits less important in the parasitic lifestyle (Jongepier et al., 2021; Schrader et al., 2021; Gautam et al., 2024). These chemoreceptors detect cuticular hydrocarbons (CHCs), a complex mixture whose compositions vary qualitatively and quantitatively. CHCs are critical for social insect communication, especially in nestmate recognition, enabling the discrimination of colony members from parasitic intruders (Dani et al., 2001; Blomquist & Bagnères, 2010; Lorenzi et al., 2011). They also contribute to desiccation resistance by sealing the cuticle and supporting thermoregulation, playing an important role in climate adaptation (Menzel et al., 2018; Sprenger et al., 2019). CHC composition can change dynamically to alleviate drought stress (Sprenger et al., 2018) or to adapt to fluctuating temperatures (Wagner et al., 1998, 2001; Martin & Drijfhout, 2009; Menzel et al., 2018; Sprenger et al., 2019), but may consequently impair detection efficiency (Wittke et al., 2022), hinting at a potential trade-off in fulfilling both functions. This highlights the challenge of optimising enemy recognition and desiccation resistance, especially in species interactions across diverse climatic landscapes.

The myrmicine ant *Temnothorax americanus* is a dulotic social parasite, which raids *Temnothorax longispinosus* colonies to capture worker brood and exploits them for their social behaviours (Wesson, 1939; Hölldobler & Wilson, 1990; Buschinger, 2009; Schmid-Hempel, 2019; Rabeling, 2021). Both species are widely distributed across northeastern North America, with parasites more abundant in southwestern regions (Jongepier et al., 2014; Macit et al., 2024). Geographical variations in parasite prevalence and thus selection pressures on local hosts prompted the evolution of divergent coevolutionary strategies. These include modifications in the CHC profile composition: *T. americanus* employs chemical insignificance by reducing its amount of recognition cues to evade host detection (Lenoir et al., 2001; Kleeberg et al., 2017; Kaur et al., 2019), while its host diversifies its colony-specific CHC profiles in areas where parasites are present, impairing advances at parasite chemical matching with hosts (Jongepier & Foitzik, 2016; Kleeberg et al., 2017; Collin et al., in review). Behavioural strategies in the host exhibit spatial shifts associated with variation in parasite abundance and climate (Jongepier et al., 2014; Segev et al., 2017; Collin et al., in review): In heavily parasitised, warm regions, hosts tend to show reduced aggression towards highly aggressive parasites, whereas in colder regions with low parasite prevalence, hosts display greater aggression towards their less aggressive sympatric parasites (Segev et al., 2017; Collin et al., in review). Following such parasite encounters, hosts temporarily elevate aggression levels and increase venom production (Pamminger et al., 2011; Scharf et al., 2011; Koenig & Moreau, 2024a), possibly in preparation for further attacks. Accordingly, hosts modify their brain transcriptome activity in response to encounters with aggressive parasites, while parasites show dynamic changes in their gene expression before and during attacks, corresponding to shifts in activity levels (Alleman et al., 2018; Kaur et al., 2019). Adding a spatial component, a previous Pool-seq study on the host *T. longispinosus* showed the antennal transcriptome strongly linked to local parasite prevalence (Macit et al., 2024). This study further provided first insights on the genomic basis of population-level host adaptation, identifying strong selection on immune functions and olfactory perception, namely selection on *peptidoglycan recognition protein* and various *odorant receptor* genes. Fundamentally, the previously unknown link between parasite prevalence and local climate conditions may crucially shape the geographic mosaic of coevolution across their heterogeneous habitats.

Here, we present the first study on the genomic basis of co-adaptation in a host and its social parasite by disentangling the effects of biotic and abiotic interactions across populations. We re-sequenced individual ants from over 120 colonies, each of the host *Temnothorax longispinosus* and its parasite *T. americanus*, across ten sites within their shared range in the Northeastern United States. For both species, we investigated: (i) population structure, (ii) genomic differentiations between populations, (iii) genome-wide associations with abiotic and biotic environmental variables (parasite prevalence and climate), (iv) genome-wide associations with key traits relevant to host–parasite interactions (chemical profiles, attack- and defence-related behaviours) obtained from our twin study (Collin et al., in review), and (v) gene expression patterns in head and fat body associated with parasite prevalence and climate. We predict that genes critical in the coevolution of these species are involved in enemy recognition (e.g., odorant receptors), signalling (e.g., cuticular hydrocarbons via fatty acid synthesis), behaviour (e.g., biogenic amines), and immune response (e.g., peptidoglycan recognition proteins). Our integrative approach establishes links between key phenotypic traits and their underlying genomic architecture, providing insights into the complex spatial dynamics of biotic and abiotic factors that shape coevolutionary patterns between social hosts and their parasites.

## Results

### Population Structure

Fifteen workers from independent colonies of the host *Temnothorax longispinosus* and between 5 and 24 individuals of the parasite *Temnothorax americanus* per population (depending on availability; Suppl. S1) were individually re-sequenced across ten populations (Fig. S1). Population structure in the host species was nearly absent, with very low pairwise FST values (mean FST = 0.021). In contrast, the parasite exhibited low but more than twice as high FST values (mean FST = 0.055; Fig. 1C). Two genotype clusters were identified in the host, whereas four distinct clusters were detected in the parasite (Fig. 1B). Isolation-by-distance was stronger in the parasite (r = 0.48, p = 0.0023) than in the host (r = 0.18, p = 0.17; Fig. 1D). Additional details are provided in the SI.

**Figure 1.**
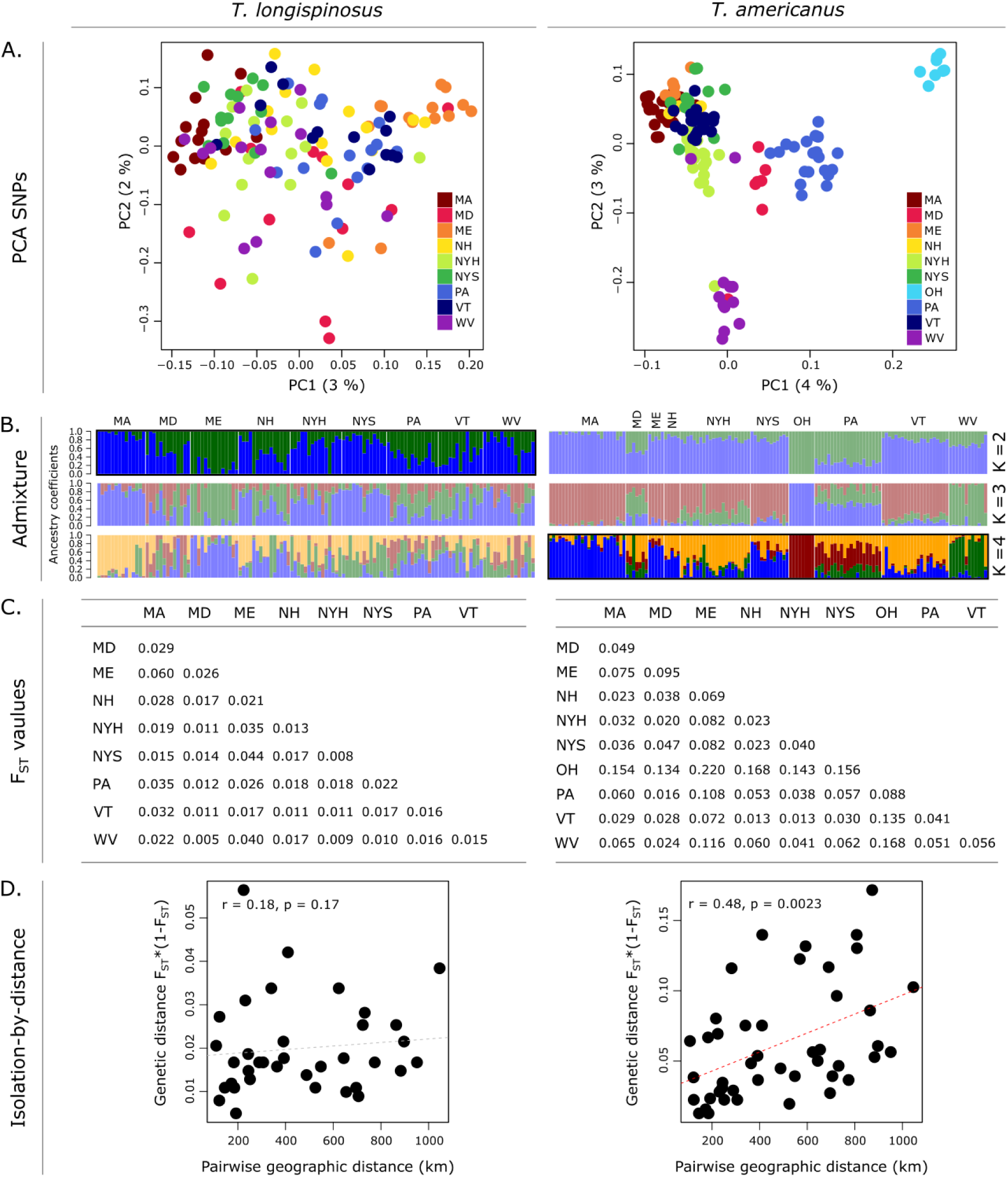
Population Structure Analysis. (A) Principal Component Analysis of SNP data, based on a thinned dataset containing ∼50k SNPs per species. (B) Admixture plot depicting the identified genotypes and their proportions for each population. Highlighted are admixture subplots for the lowest cross-entropy identified using *LEA()* ran 50x (Fig. S2A). (C) Pairwise FST values. (D) Isolation-by-distance.

### Genomic Variation Linked to Population Differentiation

We used *OutFLANK* (Whitlock & Lotterhos, 2015) to identify differentiated loci (p ≤ 0.05) based on FST outliers, indicative of local differentiations between populations. We detected 6,018 outlier loci in the host, including 282 non-synonymous SNPs (ns-SNPs) across 461 genes (Fig. 2A-B; Table S3). In contrast, the parasite exhibited only half as many outlier SNPs (3,250), but the number of ns-SNPs (177) and associated genes (405) did not differ from those of its host (padjust > 0.05 for both; Fig. S4A, S5A). Candidate genes in the host included *insulin-degrading enzyme* (*IDE*), *chitinase 10* (*CHT10*), and several *peptidoglycan recognition protein* (*PGRP*) genes (Table S4). In the parasite, notable genes included *cubilin* and *vitellogenin 1-like* and genes involved in cuticular hydrocarbon (CHC) synthesis, such as *fatty acyl-CoA reductase* and *desaturases* (Table S5). Among the 12 enriched biological functions of host genes (Fisher’s exact test, p ≤ 0.05) were immune-related processes, particularly ‘peptidoglycan catabolic process’. In the parasite, 20 enriched biological functions of parasite genes included pathways related to lipid metabolism, such as ‘lipid metabolic process’ (Fig. 2C, Suppl. S2).

**Figure 2.**
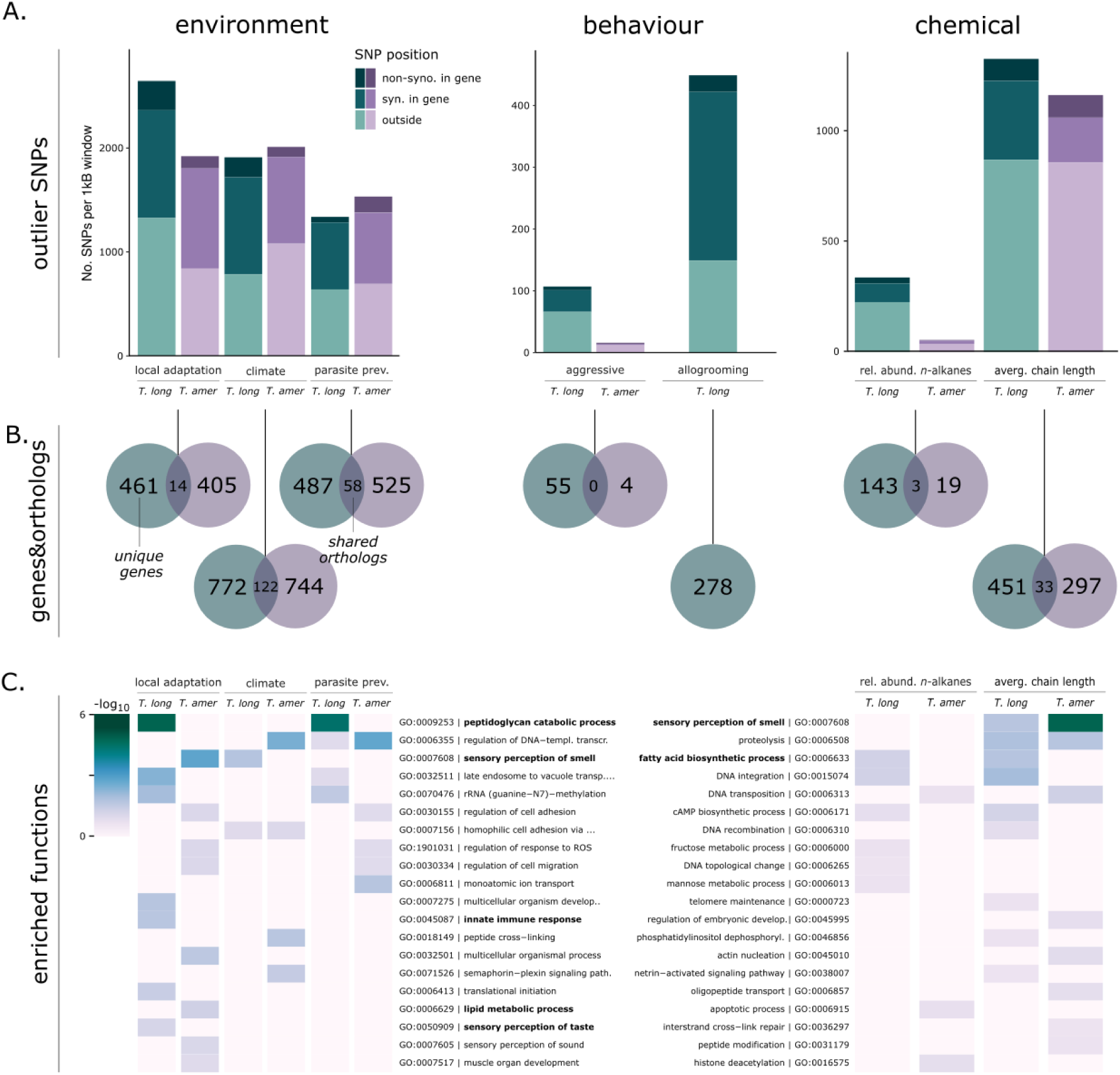
Genome-wide association analyses. (A) Total number of unlinked SNPs (i.e., 1 SNP per 1kB windows) identified by the different analyses (‘local adaptation’ refers to loci differentiated between populations), divided into non-synonymous SNPs, syn. in-exon SNPs and SNPs outside of gene regions (see Fig. S4A for more information). (B) Number of unique genes in which these SNPs reside, with overlap indicating orthologous SNP-containing genes in both species (see Fig. S5B for more information). (C) Heatmap of a collection of enriched biological GO-terms with candidate functions in bold.

### Genomic Variation Linked to Environmental Traits

We used *BayPass* v2.2 (Gautier, 2015) with its covariate model to associate genotypes with environmental factors (parasite prevalence and climate) and phenotypic traits (chemical profile and behaviour) in the host and the parasite (summary in Table S3, raw data in Suppl. S2).

### Parasite Prevalence

Parasite prevalence, defined as the relative abundance of parasites in relation to host colonies, is a proxy for parasite success and the selective pressure parasites impose on local host populations. In the host, we identified 2,706 significant SNP loci (with Bayes Factor (BF) ≥ 15) associated with population-wide parasite prevalence (including 155 ns-SNPs) across 487 genes. In contrast, we identified fewer SNPs in the parasite (1,690 SNPs, including 54 ns-SNPs), but those were distributed across more genes (525; gene count: χ² = 10.66, padjust = 0.0019; Fig. 2A-B; Fig. S5A). The number of ns-SNPs was higher in the host, as were their Bayes Factors, indicating a stronger genomic association with parasite prevalence (ns-SNPs count: χ² = 15.02, padjust < 0.001; BF: ANOVA F = 6.58, padjust < 0.03; Fig. S4A-B). Host candidate genes were similar to those identified to be differentiated between populations, including *PGRP* genes (highest BF > 50) and multiple *CHT10* genes (highest BF = 39; Fig. 3A; Table S4). These genes were also identified in a previous host Pool-seq GWAS analysis (Macit et al., 2024; Fig. S6), further supporting their role in parasite resistance/tolerance. We found that heterozygosity of significant loci within *PGRPs* was mainly differentiated between populations (ANOVA F = 2.65, p < 0.007; Fig. S3). In contrast, parasite genes associated with parasite prevalence showed little overlap with those identified to be differentiated among populations. Among the ones with the strongest signals were two *guanine nucleotide-binding protein-like 3* genes (*GNL3L*; highest BF > 50), two *multidrug resistance-associated proteins 4* (*MRP4*; highest BF > 40) and a *fatty acyl-CoA reductase* (*FAR*; highest BF = 21; Fig. 3A; Table S5). Enriched functions differed between host and parasite: In the host, again, immune-related pathways like ‘peptidoglycan catabolic process’ were among the five enriched functions, while among the five enriched functions in the parasite were those relating to transcriptional regulation, such as ‘regulation of DNA-templated transcription’ (Fig. 2C, Suppl. S2).

**Figure 3.**
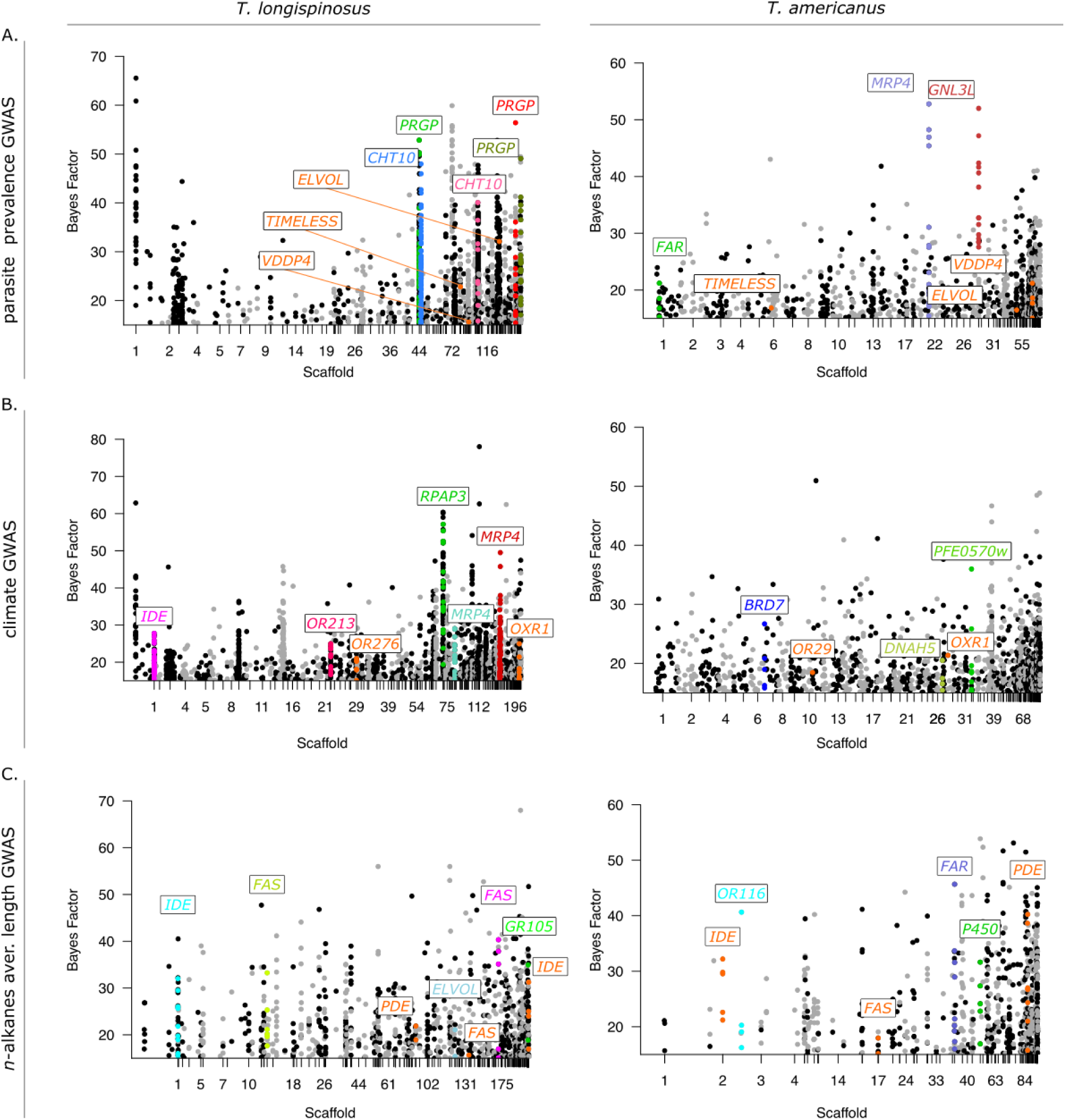
Genome-Wide Association Studies (GWAS) of *Temnothorax longispinosus* and its parasite *T. americanus*. Manhattan plots display significant single-nucleotide polymorphisms (SNPs) associated with (A) parasite prevalence, (B) climate, and (C) average chain length of *n*- alkanes. Only SNPs exceeding a Bayes Factor (BF) of 15 and located within or proximal (±2kB) to annotated genes are shown. Orthologous candidate genes between the two species are highlighted in orange (Table S6). Candidate genes presented here represent a subset of those exhibiting a high number of non-synonymous SNPs with high BF values. More details can be found in Table S4 and S5.

### Climate

By using population-specific climate data encompassing various temperature and precipitation variables (see Macit et al., 2024), we identified 3,016 SNPs associated with climate in the host (including 192 ns-SNPs) across 772 genes (Fig. 2A-B). In the parasite, we identified a similar number of associated SNPs (2,582) and genes (774) but fewer ns-SNPs (97; ns-SNP count: χ² = 9.94, padjust < 0.003; gene count: padjust > 0.05; Fig. S5A, S4A). Host genes with the strongest genomic associations included two *multidrug resistance proteins* (*MRP4*; highest BF > 40), a *trichohyalin-like* gene (highest BF > 50), and an *odorant receptor* gene (*OR*; highest BF = 23.9; Fig. 3B, Table S4). In contrast, parasite candidate genes showed significantly weaker associations than their host (BF: ANOVA F = 20.41, padjust < 0.001; Fig. S4B). Among the parasite genes with the strongest associations were *MATH and LRR domain-containing protein (PFE0570w;* highest BF = 36.0), *x-ray repair cross-complementing protein 5* (highest BF = 27.5), and a *fatty-acid reductase* (*FAR*; highest BF = 21.3; Fig. 3B, Table S5). Host climate-associated genes were enriched for five biological functions, primarily related to gene regulation and post-translational modifications (‘regulation of DNA-templated transcription’, ‘protein phosphorylation’). In contrast, the parasite’s 14 enriched functions were distinct from its host’s, with several related to metabolic and catabolic processes and signalling pathways (‘semaphorin-plexin signalling pathway’; Fig. 2C).

### Co-Adaptation

We used *OrthoFinder* v.2.5.4 (Emms & Kelly, 2015) to identify shared genes that were differentiated among populations, climate-adapted or correspond to their coevolution in both species. We identified 14 orthologous genes differentiated between populations in the host and the parasite, including *pumilio homolog 2*, linked to spatial memory in *Mus musculus* (Siemen et al., 2011) and *retinol-binding protein pinta-like*, related to visual responses in *Drosophila* (Wang & Montell, 2005; Table S6). However, the number of shared genes was not higher than expected by chance, indicating population differentiation acting on different genes and pathways in both species (hypergeometric model p = 0.22; Jaccard index = 0.021, Fig. S5B). In contrast, for both parasite prevalence and climate, the number of orthologous candidate genes identified in the GWAS was greater than expected by chance, indicating selection on the same genes (both p < 0.0001; Jaccard indices: climate = 0.10, parasite prevalence = 0.068). In the GWAS on parasite prevalence, 58 orthologs were found under reciprocal selection in both species. Many of these genes were associated with neural, synaptic, and developmental functions (*neurotrimin*, *semaphorin-1A and -2A*, *fasciclin-1, latrophilin/cirl, netrin-A*), lipid modifications and homeostasis (*elongation of very long chain fatty acids protein*, *phosphodiesterase*), circadian rhythm (*timeless*), and venom production (*venom dipeptidyl peptidase 4*; Fig. 3A and Table S6). In the climate GWAS, we identified 122 orthologous genes associated with climate adaptation in both species, twice as many as in the parasite prevalence GWAS. Several of these genes were associated with functions related to neural and photosensory perception (*fascicilin-1*, *semaphorin-2A*, *phospholipase A1*), but also to oxidative and environmental stress management (*oxidation resistance protein 1*, *carboxylic ester hydrolase*), and water retention (*nephrin*; Fig. 3B and Table S6).

### Genomic Variations Linked to Behaviour

Interactions between dulotic parasites and their hosts are mediated via behavioural traits, particularly aggression during raids. As hosts typically recognise parasites by their chemical profile, encounters often escalate to overt aggression due to chemical mismatches, suggesting targeted aggressive responses are subject to selection. Here, we use the number of aggressions after introducing an individual parasite into the host nest as a behavioural proxy (Collin et al., in review). We detected only a weak genomic signal associated with aggressive behaviour in both species. For the host, we identified 131 associated SNPs (including six ns- SNPs) across 55 genes, which included *protein groucho* (BF = 22.7), *calpain-D* (BF = 16.5), and also *insulin degrading enzyme* with several intron SNPs (highest BF = 15.8; Table S4). Those genes were enriched in four biological functions, such as neural repair and perception (‘response to axon injury’, ‘visual perception’). For the parasite, we identified only 15 SNPs (including two ns-SNPs; Table S5) within four genes, significantly less than host candidate genes (gene count: χ2 = 36.42, padjust < 0.001; Fig. 5A), which included a *juvenile hormone esterase* (BF = 15.6). Due to the low number of genes, no enriched functions could be determined (Table S5, Suppl. S2).

Host workers are often injured during raids, with their nestmates typically reacting by wound grooming. To simulate parasite-induced injuries, we observed nestmate responses to leg removal in host colonies, hypothesising grooming behaviour to be linked to parasite prevalence (see SI). Instead, we found a stronger association with local climate, with higher allogrooming frequencies in colonies from warmer regions (χ² = 7.36; p = 0.007; Fig. S14), which we explained as preventing bacterial proliferation in warmer temperatures. We identified a strong genomic basis for this behaviour, with 523 associated SNPs (including 27 ns- SNPs) across 278 genes (Suppl. S2). Candidate genes included *neprilysin-4* (BF = 24.3) and *DNA topoisomerase 3-alpha-like* (BF = 19.5; Table S4). Among the twelve enriched biological functions were some involved in transcription and expression (‘regulation of DNA-templated transcription’, ‘regulation of gene expression’; Table S4).

### Genomic Variations Linked to the Cuticular Hydrocarbon Profile

Chemical traits play a key role in host–parasite interactions by signalling and recognition mediation via methyl-branched alkanes, as well as climate adaptations by protection against dehydration via linear *n*-alkanes. As cuticular hydrocarbon biosynthesis is mediated via a conserved pathway, similar genes may be subject to selection in both species. However, we found no associated genetic variants in either species with recognition cues identified by Collin et al. (in review). Contrarily, for the relative abundance of linear *n*-alkanes, we identified 424 associated SNPs in the host (including 27 ns-SNPs) across 143 genes, and a considerably weaker genomic basis in the parasite, with only 56 SNPs (including 3 ns-SNPs) across 19 genes. While the host possesses more ns-SNPs, their association values did not significantly differ between species (ns-SNP count: χ2 = 14.04, padjust < 0.0005; BF: ANOVA padjust = 0.52; Fig. S4A-B). Candidate genes in the host included some known to be involved in CHC biosynthesis, such as *fatty acid synthases* (*FAS*, highest BF > 40) and *cytochrome P450* genes (hereafter referred to as *P450*; highest BF = 39), along with olfactory perception genes like an *OR* (highest BF = 16) and a *gustatory receptor* gene (*GR*; BF = 19; Table S4). The parasite exhibited fewer associated genes (χ2 = 79.03, padjust < 0.001; Fig. S5A), including a *FAS* (BF = 25.5) and a *P450* gene (BF = 15.9), but no perception genes (Table S5). Among the eight enriched functions of host candidate genes were processes related to CHC biosynthesis, and meta- and catabolic processes (‘fatty acid biosynthetic process’, ‘fructose/mannose metabolic process’). The parasite had half as many enriched functions, including those relating to gene regulation and cell death (‘histone deacetylation’, ‘apoptotic process’). Two orthologs were found between the species, of which one was annotated as *cullin-3* (Table S6).

The average chain length of linear *n*-alkanes can further influence desiccation resistance properties. We found that the average chain lengths of linear *n*-alkanes differed among populations in both species (host: ANOVA F = 3.50, parasite: F = 3.67, both p < 0.01) but also between species, with the host having longer linear *n*-alkanes (Mann-Whitney U test, z = 13.62, p < 0.00001; Fig. S10, see SI). Using chain lengths as a parameter, we identified 2,216 SNPs (including 100 ns-SNPs) across 481 host genes in our GWAS. The parasite had a similar number of associated SNPs (2,114) as well as ns-SNPs (102) with similar association values (ns- SNP count and BF: both padjust > 0.5; Fig. S4A-B). However, the parasite had fewer candidate genes than the host (297; gene count: χ² = 15.19, padjust < 0.0005; Fig. S5A). Identified host candidate genes were those known in CHC biosynthesis, such as several *FAS* genes (highest BF > 40), a *very long chain fatty acids protein* (BF = 21.1), and again perception genes such as an *OR* (BF = 15.5) and *GR* (BF = 18.9; Fig. 3C, Table S4). The parasite showed similar genes, such as several *P450* (highest BF = 39.5), *FAS* (highest BF = 18.0), *FAR* genes (highest BF > 40), as well as several *ORs* (highest BF = 33.1; Fig. 3C, Table S5). We identified 15 orthologous candidate genes between the two species, more than expected by chance (Jaccard coefficient = 0.063, p < 0.001; Fig. S5B), which included a *FAS* and a *phosphodiesterase* gene (Table S6).

### Transcriptional Activity Associated with Parasite Prevalence

Our earlier study on host population pools revealed a strong correlation between global antennal gene expression and local parasite prevalence (Macit et al., 2024). Compared to that earlier study, which was conducted after only a few weeks of standardised ant husbandry, we kept ants used for this study for over eight months, so that workers might have emerged under standard laboratory conditions. We also used individual data from both species to examine whether transcriptional activity in the head or fat body was similarly linked with local parasite prevalence (Suppl. S1). Principal Component Analysis revealed no clustering according to populations (Fig. 4A-B), and none of the principal components, as a proxy for global gene expression, were associated with parasite prevalence in either species (Pearson’s correlation in both tissues: p > 0.05). Using parasite prevalence as a continuous variable in a *DESeq2* analysis, we identified a greater number of differentially expressed genes in the host compared to the parasite in both tissues (fat body: χ2 = 92.19, head: χ2 = 162.91, both padjust < 0.001). The expression of 462 genes in the host fat body was associated with parasite prevalence, including a *FAR* gene and a *FAS* gene (Fig. 4C). In contrast, only 214 genes in the parasite fat body transcriptome were associated with parasite prevalence, approximately half the number observed in the host. Significant genes included a *FAS*, a *circadian clock-controlled gene* (*CLOCK*) and a *retinal homeobox protein Rx2* (*RAX*; Fig. 4C). Among the 14 enriched functions of candidate genes in the host fat body were those related to meta- and catabolic processes (Suppl. S1). Fat body expressed genes in the parasites showed 22 enriched functions, primarily associated with neuronal and stress management. In the host head transcriptome, the expression of 167 genes was associated with parasite prevalence, including a *FAS* gene and an *odorant-binding protein* (*OBP*; Fig. 4D). Among the 13 enriched functions were primarily those in fatty acid and metabolic processes (Suppl. S1). Only one significant gene was found for the parasite’s head transcriptome (*fatty-acid amide hydrolase*; Fig. 4D).

**Figure 4.**
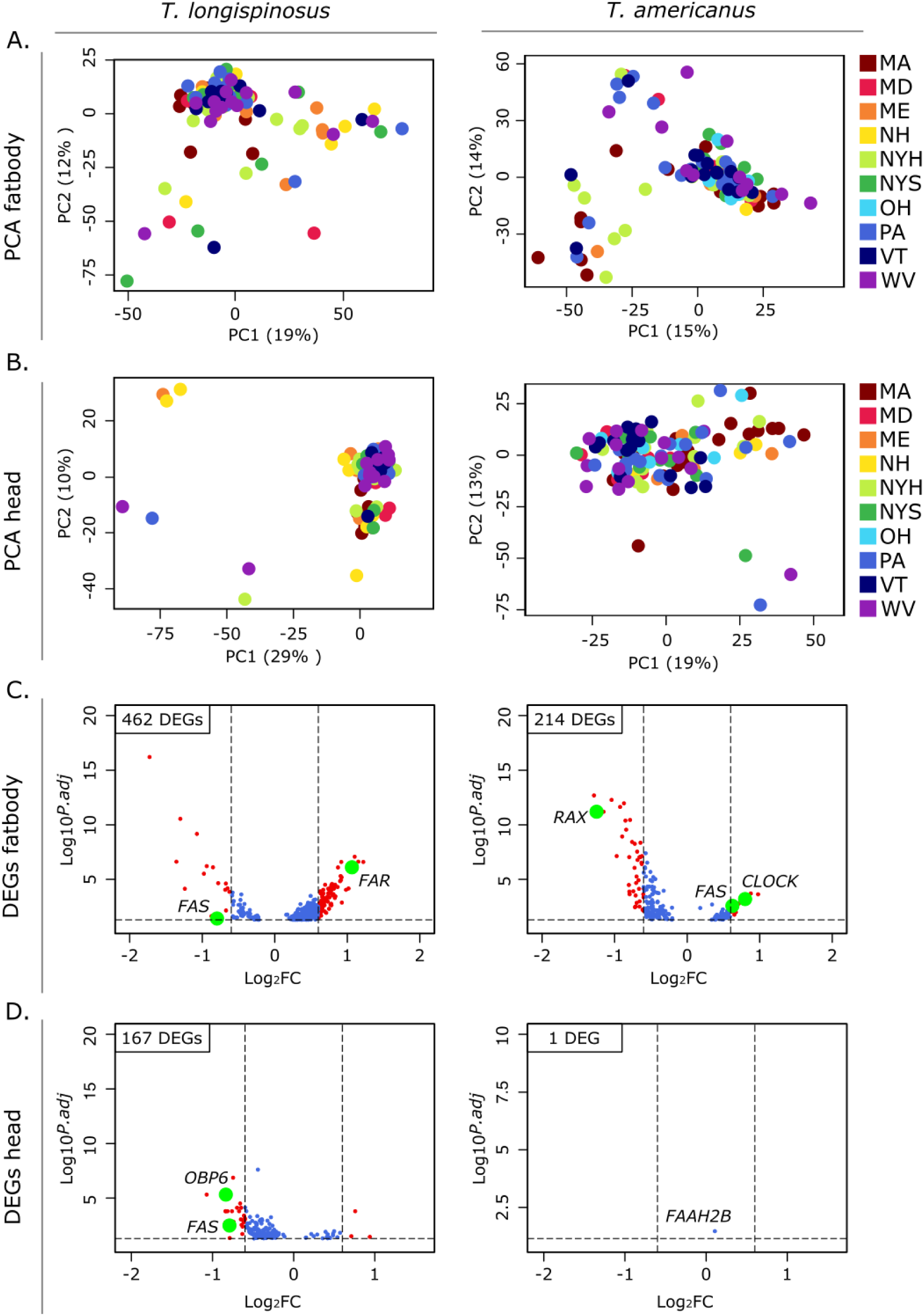
Analysis of transcriptome data. (A) PCA of fat body transcriptome and (B) head transcriptome in both species. (C) Volcano Plot of differentially expressed genes associated with parasite prevalence in the fat body and (D) in the head samples, with curated candidate genes with the highest or lowest log2FC highlighted.

Using the same transcriptome data, we analysed associations between constitutive gene expression and climate. In the fat body, more genes were climate-associated in the parasite (884) than in the host (525). Comparatively, the head transcriptome revealed fewer linked genes, with similar numbers between host (49) and parasite (40). Details, including gene functions, are provided in the supplement (Fig. S11, SI and Suppl. S1).

### Parasite-Associated Loci Linked to Differential Gene Expression

Variants within non-coding regions can be important for adaptation if they lead to altered expression of the associated genes (Nourmohammad et al., 2017). We employed an eQTL-like strategy to link SNPs identified in the parasite prevalence GWAS within regulatory regions (i.e. introns or ±2kB upstream, referred to as 2kB-SNPs) to regulatory changes associated with parasite prevalence. We identified two genes, *multidrug resistance protein* (*MRP4*) and *chitinase* 10 (*CHT10*), which are differentially expressed in the host fat body and head transcriptomes, respectively. *MRP4* contained 90 intron SNPs, and *CHT10* contained 68 intron SNPs and two 2kB-SNPs associated with parasite prevalence. In the parasite head transcriptome, the expression of *fatty acid amide hydrolase* (*FAAH*) associated with parasite prevalence was found to contain ten intron SNPs and one 2kB-SNP (Suppl. S1).

## Discussion

Coevolutionary divergence of key phenotypes in host–parasite interactions results from reciprocal selection pressures (Tellier et al., 2014; Kurtz et al., 2016), paired with selection pressures imposed by their shared environments. In species coinhabiting widespread heterogeneous environments, disentangling the genomic basis of the resulting mosaic of coevolution requires an integrative approach that accounts for the ecological context in which coevolution unfolds. Here, we examined the coevolutionary basis of the dulotic ant *Temnothorax americanus* and its host *T. longispinosus* across a broad climatic gradient in northeastern North America. By combining genome-wide association studies of environmental and phenotypic traits, and investigating gene expression, we explore both the genetic targets and regulatory architecture underlying host–parasite coadaptation.

### Disparate Population Structures Shape the Evolutionary Trajectories of Host and Parasite

Population genetic analyses revealed a marked contrast between the near-panmictic host and genetically structured parasite populations, consistent with previous studies based on neutral genetic markers (Brandt et al., 2007; Pennings et al., 2011; Pamminger et al., 2014; Macit et al., 2024). This mismatch supports the geographic mosaic theory of coevolution (Thompson, 2005), where spatially variable selection and gene flow shape coadaptive trajectories differentially. The parasite’s structure may further reflect recent range expansions into previously unparasitised regions (Jongepier et al., 2014; Macit et al., 2024) compounded by low dispersal and small effective population sizes, enhancing the effects of drift and local adaptation (Whitlock, 2004; Excoffier et al., 2009). The latter was indeed found in the parasite, showing stronger patterns of adaptations to local hosts, with the lack of hosts adapting to local parasites (Foitzik et al., 2001; Brandt & Foitzik, 2004; Foitzik et al., 2009).

### Parasite Prevalence Imposes Strong Selection on the Host

Genome-wide association analyses revealed strong asymmetries in genomic responses to parasite prevalence: host populations showed stronger and more coherent selection signals, consistent with stronger local selection pressures directly imposed by the parasite. Candidate genes in *T. longispinosus* showed enrichment in immune functions and structural defence functions, notably *peptidoglycan recognition protein*s (*PGRPs*), which typically initiate antimicrobial responses (Dziarski, 2004; Harris et al., 2015; Wang et al., 2019). Some *PGRPs* were also found to modulate behaviour via neuroimmune interactions using the gut-brain axis (Gonzalez-Santana & Diaz-Heijtz, 2020; Fioriti et al., 2024), and modulate egg-laying behaviour via octopaminergic neuronal circuits (Kurz et al., 2017), suggesting pleiotropic links between immunity and socially relevant traits. Such dual functionality may reflect genomic trade-offs or pleiotropic interactions between pathogen defence and behavioural plasticity, warranting further investigation on their precise mechanisms in social insects. Ongoing research suggests an expansion in *PGRP* genes in the genus *Temnothorax,* followed by a secondary reduction in their social parasites (pers. com. Lumi Viljakainen). Additional strong signs of selection on *chitinase 10*, involved in cuticle fortification, might indicate parasite-driven mechanical defences against injuries inflicted during raids (Shahabuddin & Kaslow, 1993; Qu et al., 2021; Rabadiya & Behr, 2024). In contrast, the parasite exhibited weaker and more diffuse selection signatures. Since parasite prevalence serves as a proxy for local population size, it may reflect a more complex, polygenic parameter shaped by interacting factors such as climate and demography. This complexity likely dilutes its role as a direct selective pressure (Vellend et al., 2014), leading to more diffuse associations (Abdellaoui & Verweij, 2021). Enriched functions of parasite prevalence-associated genes were related to gene regulatory function, suggesting dynamic regulatory mediation of adaptation. However, strong signals of selection were still observed in a few candidate genes, for instance, in *multidrug resistance-associated protein 4*, which was previously found to be downregulated during parasite raids (Alleman et al., 2018). As a key regulator of prostaglandin signalling, this gene was previously linked to reproductive and behavioural regulation in solitary and social insects (Stanley, 2006; Stanley & Kim, 2019; McAfee et al., 2024). As such, it may have a role in the behavioural shift from an inactive (queen-like, egg-laying) to an active (forager-like, raiding) lifestyle during raiding season (Blatrix & Herbers, 2004; Pohl & Foitzik, 2011). Another candidate gene with strong selection signals was *guanine nucleotide-binding protein-like 3*, a G-protein associated with learning, memory, and pheromone production (Guillén et al., 1990; Calkins et al., 2019; Liu et al., 2021), which may facilitate scouting and coordination during raids.

Shared signatures of selection in both species were found in genes related to mechanosensation (*latrophilin/cirl*; Scholz et al., 2015), circadian rhythms (*timeless*), and venom production. These genes may play critical roles in mediating host–parasite encounters by detecting colonies or intruders, coordinating raids and defences, and modulating aggression in response to each other. Selection on time-tracking genes may align with seasonal shifts in raiding and defensive behaviours (Brandt et al., 2005; Pamminger et al., 2011), contributing to the temporal organisation of antagonistic phenotypes. The function of venom-related genes under selection in both species may likely diverge: hosts employ venom in nest defences (Koenig & Moreau, 2024b), while parasites use glandular secretions from the Dufour’s gland in offensive behaviours like host manipulation (Brandt et al., 2005; Jongepier et al., 2015). These in parallel evolving genes with potential different functions echo the ‘toolkit hypothesis’, where coevolving partners repurpose conserved pathways to meet divergent ecological demands (Cini et al., 2015). Such functional overlaps highlight how common ancestry can constrain divergence while enabling asymmetric evolutionary outcomes tailored to specialised lifestyles, warranting further investigation into their specific role in host–parasite interactions.

### Climate Adaptation Shows Parallel Genomic Responses

As ectotherms, insects depend on physiological and biochemical adaptations to cope with climatic variation. Both species exhibit significant overlaps in climate-associated selection signatures, where shared genes and pathways mediate convergent physiological responses. These included genes with functions in water retention (*nephrin*; Solanki et al., 2021), stress responses (*oxidation resistance protein*; Wei et al., 2021), but also neuronal development (*semaphorin-2A*; Bates & Whitington, 2007), which may pleiotropically modulate desiccation tolerance and climate-sensitive behaviours (Segev et al., 2017). The intertwined relationship between climate and parasite prevalence is evident in climate-induced behavioural shifts during host–parasite encounters (Collin et al., in review), which can enhance, constrain, or redirect behavioural adaptations central to their coevolution under varying climatic conditions. Similar patterns have been observed in other systems, where abiotic conditions interact with biotic pressures to influence parasite dynamics via behavioural, ecological, or evolutionary pathways (Zamora-Vilchis et al., 2012; Møller et al., 2013; Bennett et al., 2016; Cable et al., 2017; Dziuba et al., 2023; Gray & Rabeling, 2023). Species-specific climate-adaptation patterns, however, also highlight their distinct life history demands: host-specific genes relate to cuticle maintenance and growth (*trichohyalin-like*, *insulin-degrading enzyme*; Stoppelli et al., 1988; Galagovsky et al., 2014), supporting long-term climatic resilience preferable for foraging, while parasite-specific genes are linked to development, metabolism, and CHC biosynthesis (*elongases*, *reductases*; Day et al., 2005; Sieron et al., 2019; Chougule et al., 2020; Farina et al., 2022), pointing to structural and physiological adjustments for their sedentary lifestyle.

### Behavioural Traits Show Weak Genomic Associations

Aggressive behaviour plays a central role in both host defence and parasite offence during raids, yet it exhibits few detectable genomic associations in either species. This likely reflects the complex and indirect genetic architecture of behavioural traits, which can obscure clear links in genotype-phenotype association studies (Abdellaoui & Verweij, 2021). Alternatively, gene expression plasticity may play a greater role than genomic divergence, as evident in substantial expression changes during raids in both species (Alleman et al., 2018). The *insulin- degrading enzyme* (*IDE*) gene was upregulated in hosts during raids (Alleman et al., 2018), and was also among the few aggression-related host candidate genes. Since older foragers are the primary defenders during parasite attacks (Koenig & Moreau, 2024a), and *IDE* has been implicated in age-dependent role shifts in *T. longispinosus* (Caminer et al., 2023), this gene likely links aggression to the transition to defending foragers in hosts. In contrast, among the few parasite-specific aggression genes was *juvenile hormone esterase*, linked to pheromone degradation and odour perception (Wei et al., 2021), likely enhancing sensitivity in detecting host aggression cues, but also caste-shifts (Mackert et al., 2008), similarly linking age- dependent shifts to aggression. In contrast, host-specific allogrooming of injured workers showed a strong genomic basis. This may be explained by its multifunctional role, including hygiene and social immunity, resulting in strong selection pressures that result in equally strong genomic signatures, similar to those identified in *Drosophila* (Yanagawa et al., 2020). Selection on chemosensory genes (*ORs* and *GRs*) associated with allogrooming underscores the importance of recognition of chemical signalling of (injured) nestmates (Sambandan et al., 2006; Yanagawa et al., 2010, 2014), while enrichments of regulatory functions suggest dynamic mediation of this behaviour, as also seen in honeybees (Hamiduzzaman et al., 2017).

### Genetic Basis of Chemical Signalling and Its Link to Chemosensory Perception

The cuticular hydrocarbon (CHC) profile in social insects serves dual functions in desiccation resistance and chemical communication (Sprenger et al., 2019), and is likely shaped by climate and host–parasite coevolution. The abundance of (linear) *n*-alkanes and their average chain lengths, both traits linked to desiccation resistance, showed strong genomic bases in both species, hinting these to be complex polygenic traits. Similar genes and functions were selected for in both species, including *fatty acid synthases*, implicated in the CHC biosynthesis, highlighting the use of convergent genetic pathways. The selection of chemosensory genes such as *ORs* and *GRs* in both species highlights the intricate pleiotropic relationship between chemical signalling and odorant perception. While previous *Hymenopteran* CHC genetic studies did not identify SNPs within perception genes (Buellesbach et al., 2022; Cohen et al., 2023), analyses on desiccation tolerance in *Drosophila* similarly found variants within *OR* and *GR* genes (Griffin et al., 2017; Rajpurohit et al., 2018). This tight evolutionary coupling of perception and signalling in both species could support a signalling function for linear *n*-alkanes previously identified in social wasps (Michelutti et al., 2018).

Using population-level data, we previously identified genomic loci associated with putative recognition cues in the host (Macit et al., 2024). However, we could not reproduce these results using colony-level data for both species, potentially due to high variability in individual recognition cues and limited statistical power from low sample sizes. popGWAS approaches (Pfenninger, in press) could offer improved power for detecting genomic associations in such variable traits and should be considered in future studies.

### Transcriptional Activity and Gene Regulation in Host–parasite Interactions

Host defence and parasite raiding behaviours are accompanied by complex transcriptional changes (Alleman et al., 2018; Kaur et al., 2019). To assess population-level variation in constitutive gene expression, we analysed transcriptomes from both species’ fat body and head tissues after eight months of standardised laboratory conditions. While this design reduces environmental noise and allows for controlled comparisons, it captures baseline, adaptive expression and may overlook important context-dependent regulatory responses as were identified in previous studies (Alleman et al., 2018; Kaur et al., 2019). Thus, the presented data reflect evolved, genetically encoded expression patterns rather than acute, plastic responses to parasites. The main ecological driver of population-level differences in constitutive gene expression differed between species: in the host, more differentially expressed genes were associated with local parasite prevalence, while in the parasite, more were associated with climate. In *T. longispinosus*, this pattern likely reflects regulatory adaptation to sustained parasite pressure. Transcriptomic analyses of pooled antennae samples showed gene expression strongly correlated with parasite prevalence but not climate (Macit et al., 2024). This is especially relevant since constitutive gene expression is costly and usually evolutionarily constrained (Wagner, 2007). This may hint at the tremendous selective pressure of parasite prevalence on this tissue, where directional selection of genotypes responsible for this beneficial gene expression pattern leads to rapid fixation within highly parasitised populations (Ghalambor et al., 2015; Campbell-Staton et al., 2017; Rivera et al., 2021). Host expressed genes associated with parasite prevalence included *odorant-binding protein* and *fatty acid synthase*, involved in chemical perception and CHC biosynthesis, linking transcriptional variation directly to recognition and signalling, both relevant in host–parasite interactions. In the parasite, the lower number of expressed genes associated with parasite prevalence may reflect both its sedentary lifestyle and simpler CHC profile (Kleeberg et al., 2017; Collin et al., in review), but also the fact that local prevalence likely reflects a weaker selective force on the parasite than on its host. Among these few differentially expressed genes associated with parasite prevalence were *FAAH*, involved in lipid signalling and neural plasticity (Ueda et al., 2000; McKinney & Cravatt, 2005; Sang & Chen, 2006), a circadian regulator linked to time-sensitive behavioural cycles, and *retinal homeobox protein*, implicated in olfactory learning (Belyaeva et al., 2009; Hofmann et al., 2016). These genes suggest gene regulation in the parasite to be more dynamic, context-dependent, and likely activated during raids. This highlights the role of timed gene expression, as evident in the importance of time-tracking genes in social parasite evolution (Feldmeyer et al., 2017), and points to flexible rather than constitutive regulatory strategies in the parasite.

## Conclusion

This study demonstrates how a social ant parasite and its host, despite shared ancestry and ecological overlap, follow divergent genomic trajectories of coadaptation across multiple populations spanning a broad interaction range. Contrasting population structures, near-panmixia in the host and pronounced genetic structuring in the parasite, generate a genomic mosaic in which coevolutionary dynamics and the degree of reciprocity vary across space. While both species exhibited similar genomic responses to prevailing climate, suggesting convergent physiological adaptation to shared environmental conditions, selection signatures linked to their coevolution and traits pivotal in their interactions were largely species-specific and reflective of their distinct lifestyle. The parasite generally showed weaker genomic signals, primarily in genes associated with behavioural shifts underlying its raiding phenotype, whereas the host displayed strong selection on immune-related genes, potentially linked to defences against parasite-induced injuries and pleiotropically the modulation of social behaviours. The difference in the strength of selection signals on the same parameter likely reflects that parasite prevalence acts as a direct selective pressure on the host, but due to its strong link to climate, presents a multilayered selective pressure for the parasite, resulting in more diffuse genetic signals. By integrating genomic, phenotypic, and environmental data, this study unravels how coadaptation arises from the intricate interplay of population structure, reciprocal selective pressures, and ecological context, underscoring the highly context- and role-specific nature of social parasitism’s evolution in heterogeneous landscapes.

## Materials and Methods

### Sample Collection and Estimation of Parasite Prevalence

Colonies of *Temnothorax longispinosus* and its social parasite, *T. americanus*, were collected from ten locations across the northeastern USA (Fig. S1, Table S1 and S2). Sampling occurred near roads and tracks in state parks and on private property, with permission obtained. Ants were brought to Mainz, Germany, and maintained under standard laboratory conditions for eight months before dissection. The accompanying behavioural and chemical data from Collin et al. (in review) were obtained four additional months later. Parasite prevalence, calculated as the percentage of parasite colonies within the local *Temnothorax* community, was determined using long-term collection data (Herbers & Foitzik, 2002; Brandt & Foitzik, 2004; Achenbach & Foitzik, 2009; Jongepier et al., 2014; Kaur et al., 2019; Macit et al., 2024).

### Sample Preparation, Sequencing and Pre-Processing

Fifteen independent colonies per population and species were sampled (host: min = 14 max/mean/median = 15; parasite: min = 5, max = 24, mean = 15, median = 18). DNA from the thorax, and RNA from the fat body and head were extracted. Whole-genome sequencing (WGS) and RNA sequencing (RNA-seq) were performed on an Illumina NovaSeq 6000 platform by Novogene. Both were quality-checked with *FastQC* v.0.11.9 (Andrews et al., 2015) and trimmed using *Trimmomatic* v.0.39 (Bolger et al., 2014). WGS data were mapped using *BWA mem* v.0.7.17 (Li et al., 2009), and RNA-seq data were mapped using *HISAT2* v.2.1.0 (Kim et al., 2015) to unpublished reference genomes (Boomsma et al., 2017). For WGS data, variants were called using *BCFtools* v1.16 (Li, 2011) using predetermined filtering parameters, resulting in similar numbers of SNPs in both species (host: 1,677,757 SNPs; parasite: 1,604,099 SNPs). Transcript read count tables were generated using *HTSeq* v.2.0.2 (Putri et al., 2022) from RNA-seq data, and used for *DESeq2* v.1.42.0 (Love et al., 2014) in *R* v.4.3.2 (R Core Team, 2021) using parasite prevalence as a continuous variable. The analysis was repeated using climate PC1 values (obtained from Macit et al., 2024) as a continuous variable and will be reported on in the SI. Further information on candidate genes in both WGS and RNA-seq data was collected using *InterPro* v.5.61.93 (Paysan-Lafosse et al., 2023), retrieving Gene Ontologies (GO) to perform *topGO* v.2.54.0 (Alexa & Rahnenfuhrer, 2017) and *BlastP* v.2.13.0 (Altschul et al., 1990), searching against the non-redundant invertebrate database (retrieved on NBCI on Jan 2022), and proteomes of *Drosophila melanogaster* and *Apis mellifera* (retrieved on UniProt on Jan 2024; Proteome IDs: UP000000803 and UP000005203; UniProt Consortium, 2018). Any gene names given in this study refer to the best blast hit (based on e-value) generated in *A. mellifera* if not stated otherwise. Orthologs were identified using *OrthoFinder* v.2.5.4 (Emms & Kelly, 2015). Chi-square tests in *R* assessed significant differences in candidate gene numbers (gene numbers: *T. longispinosus*: 16,064, *T. americanus*: 14,128) and ns-SNPs (exon lengths: *T. longispinosus*: 21,924,724, *T. americanus*: 20,133,265). Bayes Factors of ns-SNPs were log-transformed and analysed for differences between species using a one-way ANOVA in *R*. Significant overlaps in orthologous candidate genes were determined using the *hypergamous()* function from *SciPy* (Virtanen et al., 2020). All statistical tests used a significance threshold of FDR-corrected p ≤ 0.05.

### Population Structure, Local Adaptation and GWAS

Population structure was analysed using the *R* package *sambaR* (De Jong et al., 2021), incorporating the *findstructure()* function. The lowest cross-entropy was observed at k = 2 for the host and k = 4 for the parasite (Fig. S2A). Locally differentiated SNPs (p ≤ 0.05) were identified using *OutFLANK* (Whitlock & Lotterhos, 2015) within the *selectionanalyses()* function in *sambaR*. Genome-wide association studies (GWASs) were conducted using *BayPass* v2.2 (Gautier, 2015) in its standard covariate mode. The parameters employed to identify their genomic basis included: (i) parasite prevalence, (ii) climate parameters, (iii) chemical profiles (relative abundance of recognition cues, (linear) *n*-alkanes, and their average chain length) and (iv) aggressive behaviour, important for both parasitic raids and host defences, and allogrooming for the host. Other statistically significant species-specific behaviours, such as host brood-carrying and parasite passive behaviour, were similarly analysed but presented in the SI only. Data for chemical and behavioural analyses largely originated from Collin et al. (in review) from sister ants genotyped in this study. Identified SNPs were considered significant at a Bayes Factor of ≥ 15. A detailed description of the methods used in this study can be found in the SI.

## Supporting information

Supplement S1 - Sample Info and RNA-seq Results

Supplement S2 - GWAS Results

Supplement S3 - Raw Data Injury Assay

## Acknowledgements

We thank Jennifer Lee Grossmann, Jonas Wittig, Marcel Adrian Caminer, and Sophie Späth for their help in ant collection. We thank Dr. Menno de Jong for his help with *sambaR*, and Dr. Juliane Hartke for her help with retrieving climate data. We used ChatGPT for assistance in improving our writing in some sections.

## Author Contribution

SF and BF conceived the study. MNM, EC, SF, and BF designed the experimental setup and collected ant colonies. MNM and EC sampled colonies and performed dissections. MK performed DNA and RNA extractions. MNM performed all population genomic and transcriptomic analyses, supervised by BF and partially MP. MNM and MENB performed demographic history analysis. EJ and ML conducted injury experiments, which were analysed by EC, and supervised by SF. MNM wrote the first draft of the manuscript, and all authors revised it. The authors declare no conflict of interest.

## Funding

This study was funded by the German Research Foundation (DFG) in a grant to BF (FE 1333/3- 3) and SF (Fo298/17-3) and in GRK 2526/1—Project number 407023052. EC received funding from the Huyck Preserve, New York.

## Data Availability

The following supplementary material is available under open access: Supplement S1, S2 and S3, and Supporting Information (including Supplementary Tables, Figures, Methods, Results and additional Discussion). Raw sequence data were uploaded to the European Nucleotide Archive (ENA) and are accessible under study accession no. PRJEB76961.

## Notes

### Competing Interest Statement

The authors have declared no competing interest.

