## Supplementary Information for "The Genomic Basis of Social Parasitism: A Geographical Mosaic of Behavioural, Chemical, and Environmental Adaptations in a Widespread Host–Parasite System"

[*]shared first authors

[§]shared last authors

#### Table of Contents


| Description | | Page |
|---|---|---|
| **Table S1.** | Summary of Collection Sites | 1 |
| **Table S2.** | Detailed Location and Coordinates of all Subsites | 2-3 |
| **Table S3.** | Summary of GWASs | 4 |
| **Table S4.** | GWAS Candidate Genes in *T. longispinosus* | 5 |
| **Table S5.** | GWAS Candidate Genes in *T. americanus* | 6 |
| **Table S6.** | Orthologous Candidate Genes | 7 |
| **Figure S1.** | Collection Map | 8 |
| **Figure S2.** | Suppl. Information on Population Structure Analysis | 9 |
| **Figure S3.** | *PGRP* Genotypes | 10 |
| **Figure S4.** | Number of non-syn. SNPs and their Bayes Factors | 11 |
| **Figure S5.** | Number of Candidate Genes and Orthologs | 12 |
| **Figure S6.** | Overlaps with PoolSeq Study | 13 |
| **Materials and Methods** | | 14-24 |
| **Additional Results and Discussion** | | 25-38 |
| **Figure S7.** | Population Structure analysis including Outlier Ohio samples | 27 |
| **Figure S8.** | Nucleotide Diversity and Watterson's theta | 28 |
| **Figure S9.** | Demographic History Analysis | 30 |
| **Figure S10.** | Variations in Linear *n*-alkane Chain Lengths | 32 |
| **Figure S11.** | Volcano Plots of Transcriptome Analyses | 38 |
| **First Aid in Ants: Response of *Temnothorax* Ants to Injury in Relation to Social Parasite Prevalence and Climate** | | 49-45 |
| **Figure S12.** | Experimental Design for Injury Assays | 40 |
| **Table S7.** | Ethogram of Observed Behaviours | 42 |
| **Figure S13.** | Behavioural Responses to Injury | 43 |
| **Figure S14.** | Effects of Climate on Allogrooming Duration and Occurrence | 44 |
| **References** | | 45-52 |

**Table S1. Summary of collection sites.** States and their abbreviations, coordinates, and number of collected colonies of host and parasite and their long-term parasite prevalence as well as their prevalence based solely on collections from our preliminary study (Macit et al., 2024) (see M&M section in that study for more information about the estimates).

| Population | Location | Coord. | *T. longispinosus* | *T. americanus* | Parasite prevalence* | Parasite prevalence 2021 |
|---|---|---|---|---|---|---|
| Massachusetts (MA) | Beaver Brook North and Rock Meadow Reservation | 42.407, -71.204 | 501 | 69.86 | 12.233 | 13.024 |
| Maryland (MD) | New Germany State Park | 39.611, -79.134 | 112 | 11 | 8.943 | 8.943 |
| Maine (ME) | Bethel Community Forest | 44.413, -70.851 | 504 | 13 | 2.515 | 4.375 |
| New Hampshire (NH) | Belknap Mountain State Forest | 43.510, -71.402 | 423 | 10 | 2.309 | 8.333 |
| New York Huyck (NYH) | Huyck Preserve | 42.525, -74.160 | 5634 | 780 | 12.161 | 18.863 |
| Newy York South (NYS) | Fahnestock State Park | 41.455, -73.868 | 178 | 25.62 | 12.583 | 12.583 |
| Ohio (OH) | Private Property | 41.753, -80.967 | 573 | 105.58 | 15.558 | 13.427 |
| Pennsylvania (PA) | S.B. Elliot State Park | 41.124, -78.523 | 314 | 53 | 14.441 | 14.441 |
| Vermont (VT) | Branbury State Park | 43.945, -73.076 | 1787 | 140 | 7.265 | 10.460 |
| West Virginia (WV) | Watoga State Park | 38.110, -80.136 | 843 | 139.65 | 14.211 | 16.977 |

* long-term data from Macit et al. (2024), Kaur et al. (2019), Jongepier et al. (2014), Foitzik et al. (2009), Brandt & Foitzik (2004), and Herbers & Foitzik (2002)

**Table S2. Detailed location and coordinates of all subsites.**

| State | Subsite | Coordinates | Location |
|---|---|---|---|
| MA | a | 42.410836, -71.208085 | Beaver Brook North Reservation |
| | b1 | 42.410836, -71.208085 | Rock Meadow Conservation Area |
| | b2 | 42.399000, -71.194361 | |
| ME | a | 44.406120, -70.861389 | West Bethel Park |
| | b | 44.407185, -70.866737 | Bethel Community Forest- West |
| | c | 44.415182, -70.857712 | |
| | d | 44.421951, -70.819382 | Bethel Community Forest - East |
| MD | a | 39.629078, -79.145439 | New Germany State Park |
| | b | 39.514118, -79.156654 | |
| | c1 | 39.570572, -79.201034 | Big Run State Park |
| | c2 | 39.627922, -79.107284 | Roadside |
| | d | 39.627922, -79.107284 | Private Property* |
| | e | 39.638573, -79.105064 | Behind Trinity Cemetery |
| | f | 39.628662, -79.125107 | Acorn Loop Hiking Trail, New Germany State Park |
| | g | 39.650993, -79.122215 | Meadow Mountain Trail, New Germany State Park |
| NH | a | 43.500557, -71.413171 | Private Property* |
| | b | 43.516384, -71.379341 | Belknap Mountain State Forest |
| | c | 43.513794, -71.413536 | Private Property* |
| NYH | a1 | 42.520329, -74.146408 | Huyck Preserve |
| | a2 | 42.515518, -74.161789 | |
| | b | 42.515518, -74.161789 | |
| | c | 42.528486, -74.160333 | |
| | d | 42.532192, -74.162888 | |
| | e | 42.533791, -74.164268 | |
| | f | 42.525547, -74.170349 | |
| | g | 42.530544, -74.151466 | |
| NYS | a | 41.445290, -73.865179 | Fahnestock State Park |
| | b | 41.450997, -73.857043 | |
| | c | 41.480529, -73.917455 | Private Property* |
| | d | 41.443157, -73.867508 | Stone Garden at Fahnestock State Park |
| | e | 41.454262, -73.833420 | Tree Lake Trail at Fahnestock State Park |
| OH | a | 41.759503, -80.965593 | Private Property* |
| | b | 41.759514, -80.966522 | |
| | c | 41.758891, -80.966794 | |
| | d | 41.758312, -80.965341 | |
| | e | 41.755195, -80.946406 | |

| | | | |
|---|---|---|---|
| | f | 41.757405, -80.942399 | |
| | g | 41.749222, -80.948235 | |
| | h | 41.743347, -80.958672 | |
| | i | 41.744363, -81.031441 | |
| | j | 41.748822, -80.974843 | |
| | k | 41.747403, -80.968187 | |
| **PA** | a | 41.137976, -78.516263 | S.B. Elliot State Park |
| | b | 41.111185, -78.528163 | |
| | c | 41.102461, -78.526507 | |
| | d | 41.118119, -78.518040 | |
| | e | 41.137276, -78.520605 | |
| | f | 41.126563, -78.528959 | |
| | g | 41.136524, -78.505280 | |
| | h | 41.139987, -78.503009 | |
| | i | 41.118080, -78.516296 | |
| | j | 41.113468, -78.571419 | |
| **VT** | a1 | 43.965164, -73.076485 | Branbury State Park |
| | a2 | 43.972078, -73.072761 | |
| | b | 43.944454, -73.102104 | |
| | c | 43.972078, -73.072761 | |
| | d | 43.971760, -73.086716 | |
| | e | 43.979463, -73.064017 | |
| | f | 43.973076, -73.059197 | |
| | g | 43.969719, -73.083763 | |
| | h | 43.924728, -73.096313 | |
| | i | 43.882782, -73.063850 | |
| | j | 43.842194, -73.060585 | |
| **WV** | a | 38.124519, -80.114105 | Watoga State Park |
| | b | 38.106472, -80.129555 | Ann Bailey Trail, Watoga State Park |
| | c | 38.107899, -80.131996 | |
| | d | 38.107495, -80.134593 | |
| | e | 38.119175, -80.154663 | Watoga Park Road, Watoga State Park |
| | f | 38.102936, -80.145622 | Ann Bailey Trail, Watoga State Park |
| | g | 38.102665, -80.141525 | |
| | h | 38.106857, -80.136772 | |

* Permission granted

**Table S3. Summary of local adaptation analysis using *OutFLANK* (Whitlock & Lotterhos, 2015) and GWASs using *BayPass* (Gautier, 2015)**

| species | type of trait | type of covariant | parameter source | #samples | #SNPs (exon : intron : 2kB : outside) | #genes (% to all genes) | #enriched functions |
|---|---|---|---|---|---|---|---|
| ***T. longispinosus*** | general local adaptation | | Macit et al., 2024 | 127 | 6018 (396 : 2822 : 529 : 2271) | 461 (2.87) | 11 |
| | environment | climate | | 127 | 3016 (260 : 1287 : 395 : 1074) | 772 (4.81) | 5 |
| | | parasite prevalence | | 127 | 2706 (221 : 1184 : 289 : 1012) | 487 (3.03) | 5 |
| | behaviour | aggressive | Collin et al., in review | 96 | 131 (9 : 33 : 18 : 71) | 55 (0.34) | 4 |
| | | brood carrying | | 96 | 39 (1 : 5 : 4: 29) | 10 (0.062) | 0 |
| | | allogrooming | this study | 66 | 523 (40 : 284 : 40 : 159) | 278 (1.75) | 12 |
| | chemical | rel. abund. recog. cues | Collin et al., in review | 127 | 0 | 0 | 0 |
| | | rel. abund. linear *n*-alkanes | | 127 | 424 (31 : 91 : 55 : 247) | 143 (0.89) | 8 |
| | | aver. chain length *n*-alkanes | | 127 | 2216 (121 : 550 : 282 : 1263) | 481 (2.81) | 11 |
| ***T. americanus*** | general local adaptation | | Macit et al., 2024 | 137 | 3250 (240 : 1458 : 351 : 1201) | 405 (3.44) | 20 |
| | environment | climate | | 137 | 2582 (124 : 911 : 229 : 1318) | 744 (5.27) | 14 |
| | | parasite prevalence | | 137 | 1690 (73 : 728 : 141 : 748) | 525 (3.72) | 5 |
| | behaviour | aggressive | Collin et al., in review | 49 | 15 (2 : 2 : 0 : 11) | 4 (0.028) | 0 |
| | | passive | | 49 | 35 (0 : 5 : 6 : 24) | 11 (0.078) | 0 |
| | chemical | rel. abund. recog. cues | | 116 | 0 | 0 | 0 |
| | | rel. abund. linear *n*-alkanes | | 116 | 56 (4 : 14 : 2 : 36) | 19 (0.13) | 4 |
| | | aver. chain length *n*-alkanes | | 116 | 2114 (121 : 389 : 223 : 1381) | 297 (2.1) | 13 |

**Table S4. Selection of GWAS candidate genes for host species *Temnothorax longispinosus*.** In bold red are genes that were also identified as candidates in one of the other GWAS, and in bold orange are genes that have an orthologous candidate gene in the parasite (Table S6). Gene names with asterisks were highlighted in the Manhattan plots in Fig. 3A-C. The full list of SNPs and genes can be found in Suppl. S2.

| **genomic association** | | **candidate genes (#nsSNP, highest BF)** | **gene name** | **selection of enriched functions** |
|---|---|---|---|---|
| **local adaptation** | | **Tlon_g15805** (17) | *unc-45 homolog B* | GO:0009253 peptidoglycan catabolic process<br>GO:0045087 innate immune responses<br>GO:0005975 carbohydrate metabolic process<br>GO:0009081 branched-chain amino acid metabolic process |
| | | Tlon_g11697 (12) | *mesh* | |
| | | **Tlon_g00264** (10) | *insulin-degrading enzyme* | |
| | | **Tlon_g13447** (6) | *chitinase 10* | |
| | | Tlon_g16880 (6) | *trichohyalin-like* | |
| | | **Tlon_g13446** \| **g13570** \| **g07266** \| g07261 (17) | *PGRP* | |
| **parasite prevalence** | | **Tlon_g13446** \| **g13570** \| **g07266** \| g07269 \| g07264 \| g07263 \| g13445 (18, 52.8) | *PGRP** | GO:0009253 peptidoglycan catabolic process<br>GO:0045087 innate immune responses |
| | | **Tlon_g13447** \| g13449 \| g00818 \| g00817 (8, 39.1) | *chitinase 10 (cht10)** | |
| | | **Tlon_g15805** \| g05389 \| g15802 (8, 33.1) | *unc-45 homolog B* | |
| **climate** | | Tlon_g04123 \| g17850 (17, 45.8) | *multidrug resistance-associated protein 4 (MRP4)** | GO:0006355 regulation of DNA-templated transcription<br>GO:0006468 protein phosphorylation |
| | | Tlon_g16886 (7, 52.6) | *RNA polymerase II-associated protein 3 (RPAP3)** | |
| | | TlonOR213 \| **OR276** (4, 23.9) | *odorant receptor gene** | |
| | | Tlon_g16880 (10, 60.4) | *trichohyalin-like* | |
| | | **Tlon_g00264** (4, 20.5) | *insulin-degrading enzyme* | |
| **aggression** | | Tlon_g09388 (1, 22.7) | *protein groucho* | GO:0048678 response to axon injury<br>GO:0007601 visual perception |
| | | Tlon_g09021 (1, 16.5) | *calpain-D* | |
| | | Tlon_g11073 (1, 15.1) | *F-box/LRR-repeat protein 4 (FBXL4)* | |
| **allogrooming** | | Tlon_g04574 (1, 24.3) | *neprilysin-4* | GO:0006355 regulation of DNA-templated transcription<br>GO:0010468 regulation of gene expression |
| | | Tlon_g14038 (1, 18.5) | *RNA exonuclease 1* | |
| | | Tlon_g01170 (2, 19.5) | *DNA topoisomerase 3-alpha-like* | |
| ***n*-alkanes** | **rel. abund.** | TlonOR120 \| **OR436** (17, 22.6) | *odorant receptor* | GO:0006633 fatty acid biosynthetic process |
| | **aver. chain** | Tlon_g05168 \| g08044 \| g05010 \| g02972 (7, 40.3) | *fatty acid synthase (FAS)** | GO:0006633 fatty acid biosynthetic process<br>GO:0007608 sensory perception of smell |
| | | Tlon_g01986 (2, 21.1) | *elongation of very long chain fatty acids(ELVOL)** | |
| | | Tlon_g00265 \| **g13585** (3, 31.3) | *insulin-degrading enzyme (IDE)** | |
| | | **TlonOR436** (1,15.5) | *odorant receptor* | |
| | | TlonGR105 (1, 18.9) | *gustatory receptor (GR)** | |

**Table S5. Selection of GWAS candidate genes for the parasite *Temnothorax americanus*.** In bold red are genes that were also identified as candidates in one of the other GWAS, and in bold orange are genes that have an orthologous candidate gene in the host (Table S6). Gene names with asterisks were highlighted in the Manhattan plots in Fig. 3A-C. The full list of SNPs and genes can be found in Suppl. S2

| genomic association | | candidate genes (#nsSNP; highest BF) | gene name | selection of enriched functions |
|---|---|---|---|---|
| **local adaptation** | | Tame_g13193 (10) | *cubilin* | GO:0006629 lipid metabolic process<br>GO:0006869 lipid transport<br>GO:0007605 sensory perception of sound |
| | | Tame_g13414 (8) | *fatty acyl-CoA reductase* | |
| | | Tame_g09688 (5) | *cytochrome P450* | |
| | | Tame_g09621 (5) | *vitellogenin 1-like* | |
| | | Tame_g09678 (4) | *long-chain-fatty-acid--CoA ligase* | |
| | | Tame_g00717 (3) | *circadian clock-controlled protein* | |
| | | Tame_g13242 (3) | *acyl-CoA Delta(11) desaturase* | |
| **parasite prevalence** | | Tame_g08420 (4, 52.0) | *guanine nucleotide-binding protein-like 3 (GNL3L)** | GO:0006355 regulation of DNA-templated transcription<br>GO:0006370 7-methylguanosine mRNA capping |
| | | Tame_g06605 \| g06604 (4, 48.2) | *multidrug resistance-associated protein 4 (MRP4)** | |
| | | **Tame_g00154** (4, 21.2) | *fatty acyl-CoA reductase (FAR)** | |
| **climate** | | **Tame_g00154** (2, 21.3) | *fatty acyl-CoA reductase* | GO:0008152 metabolic process<br>GO:0030149 sphingolipid catabolic process<br>GO:0071526 semaphorin-plexin signalling pathway<br>GO:0007166 cell surface receptor signalling pathway |
| | | Tame_g09659 (8, 36.0) | *PFE0570w** | |
| | | Tame_g08104 (3, 20.5) | *dynein heavy chain 5 (DNAH5)** | |
| | | Tame_g13489 (2, 20.8) | *bromodomain-containing protein 7 (BRD7)** | |
| | | Tame_g16378 \| g13911 (4, 27.5) | *x-ray repair cross-complementing protein 5 \| 6* | |
| **aggression** | | Tame_g14891 (1, 15.6) | *carboxylic ester hydrolase* | |
| ***n*-alkanes** | **rel. abund.** | **Tame_g04921** (1, 25.5) | *fatty acid synthase* | GO:0016575 histone deacetylation<br>GO:0006915 apoptotic process |
| | | **Tame_g14411** (1, 15.9) | *cytochrome P450* | |
| | **aver. chain** | Tame_g11443 \| **g14411** (7, 39.5) | *cytochrome P450 (P450)** | GO:0007608 sensory perception of smell<br>GO:0006857 oligopeptide transport<br>GO:0031179 peptide modification |
| | | TameOR116 \| OR216 \| OR319 \| OR204 (5, 20.8) | *odorant receptor (OR)** | |
| | | **Tame_g06476** \| **g04921** \| g06477 (4, 18.0) | *fatty acid synthase (FAS)** | |
| | | Tame_g10337 \| g10334 (2, 45.7) | *fatty-acyl-CoA reductases (FAR)** | |

**Table S6. Orthologous candidate genes identified in GWAS of both host *T. longispinosus* and parasite *T. americanus* species.** Gene names with asterisks were highlighted in orange in the Manhattan plots in Fig. 3A-C. Details can be found in Suppl. S2.

| genomic association | | orthoGroup (Tlon \| Tame) | gene name |
|---|---|---|---|
| **local adaptation** | | OG0000567 (Tlon_g00462 \| Tame_g01233) | *cytochrome P540* |
| | | OG0001201 (Tlon_g03331 \| Tame_g03182) | *pumilio homolog 2* |
| | | OG0004851 (Tlon_g12994 \| Tame_g05034) | *heterogeneous nuclear ribonucleoprotein U* |
| | | OG0000100 (Tlon_g16890\| Tame_g08624) | *retinol-binding protein pinta-like* |
| **parasite prevalence** | | OG0001670 (Tlon_g18643 \| Tame_g15493) | *venom dipeptidyl peptidase 4 (VDP4)** |
| | | OG0000523 (Tlon_g17222 \| Tame_g13159, Tame_g13161) | *protein timeless homolog (TIMELESS)** |
| | | OG0000957 (Tlon_g04125 \| Tame_g12517) | *elongation of very long chain fatty acids protein (ELVOL)** |
| **climate** | | OG0006318 (Tlon_g09719 \| Tame_g08220) | *oxidation resistance protein 1 (OXR1)** |
| | | OG0000059 (TlonOR272,TlonOR274,TlonOR276 \| TameOR29) | *odorant receptor (OR)** |
| ***n*-alkanes** | **rel. abund.** | OG0000233 (Tlon_g13654 \| Tame_g00605) | *cullin-3* |
| | **aver. chain** | OG0000018 (Tlon_g02972\| Tame_g06476) | *fatty acid synthase (FAS)** |
| | | OG0010382 (Tlon_g18775 \| Tame_g16435) | *phosphodiesterase (PDE)** |
| | | OG0000036 (Tlon_g13585 \| Tame_g05743 ) | *insulin-degrading enzyme (IDE)** |

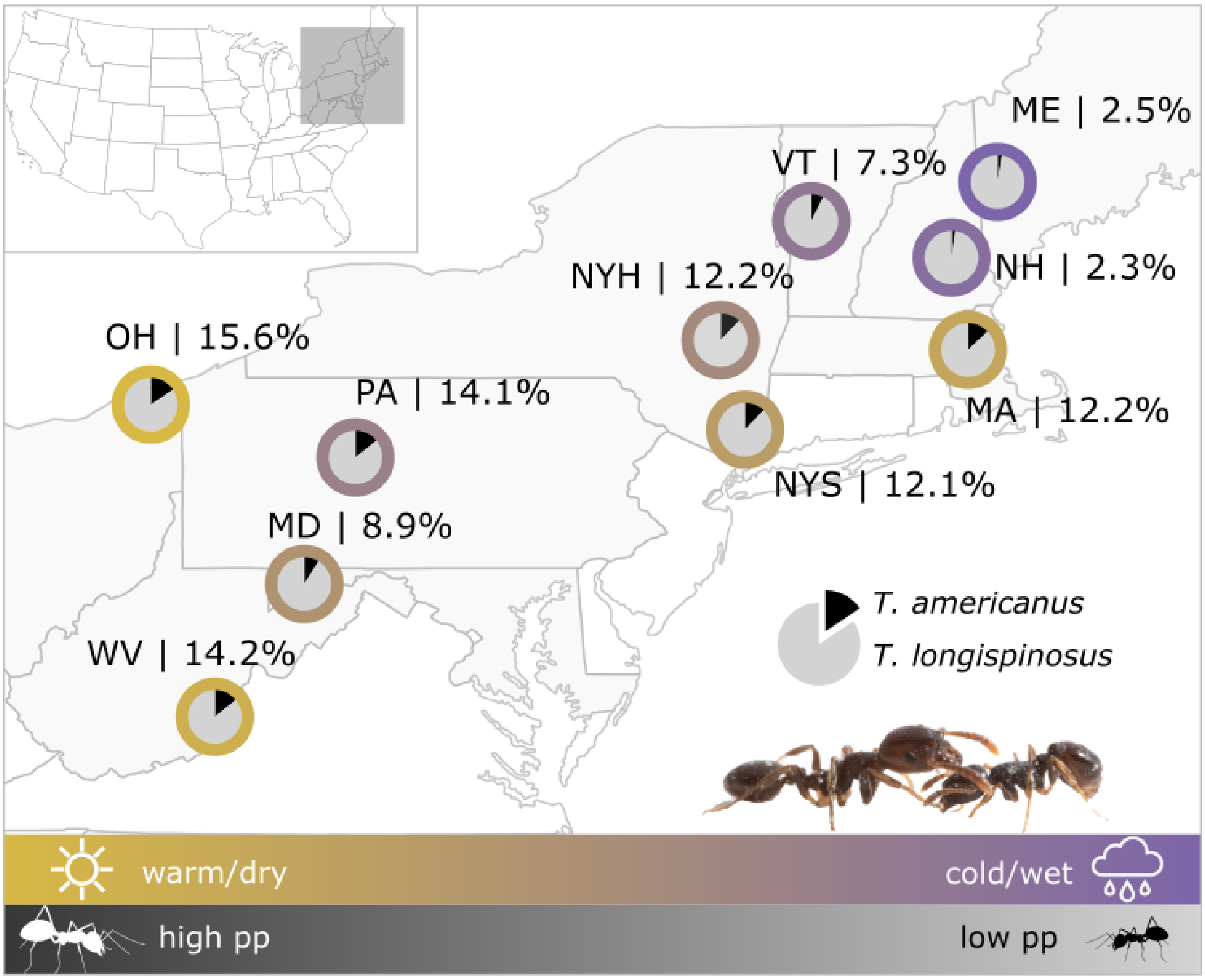


**Figure S1. Geographic distribution and sampling of *Temnothorax longispinosus* and its parasite *T. americanus* in the northeastern United States.** Sampling sites (see Tables S1 and S2 for details) are depicted using pie charts that indicate local parasite prevalence (percentages shown above) and colour gradients representing local climate conditions, ranging from warm and dry (yellow) to cold and wet (purple), as described in Macit et al. (2024). Photo © Romain Libbrecht.

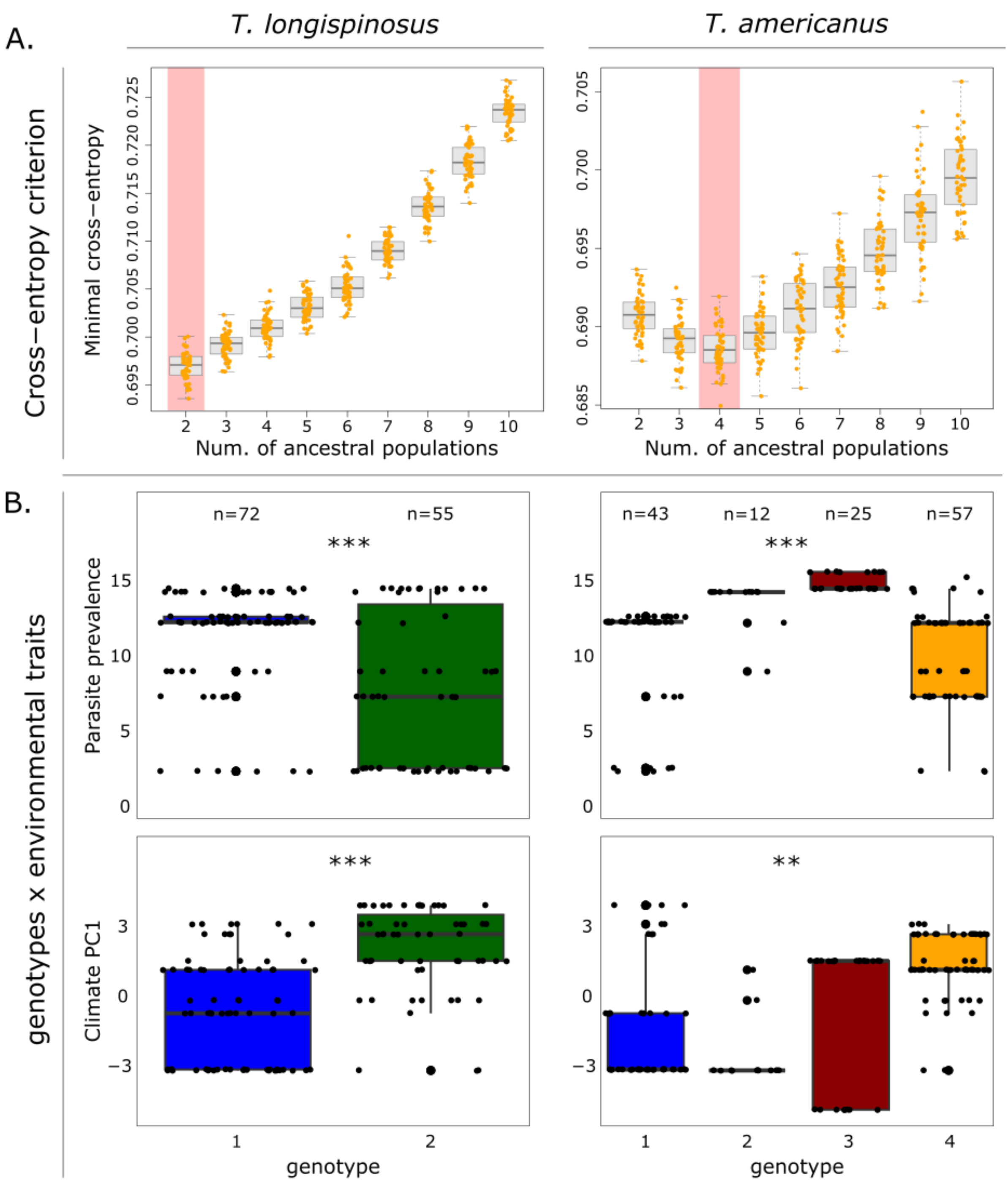


**Figure S2. Population structure and environmental associations of genotypes.** (A) Determination of the optimal number of ancestral populations (k) based on the cross-entropy criterion for k = 2 for *T. longispinosus* and k = 4 for *T. americanus*. (B) Associations of individual genotypes and environmental variables (parasite prevalence and climate PC1). ANOVA tests revealed significant effects of both parasite prevalence and climate on genotype association in *T. longispinosus* (parasite prevalence: $F = 42.72$, $p < 0.0001$; climate PC1: $F = 30.75$, $p < 0.0001$) and *T. americanus* (parasite prevalence: $F = 46.84$, $p < 0.001$; climate PC1: $F = 9.01$, $p = 0.0032$).

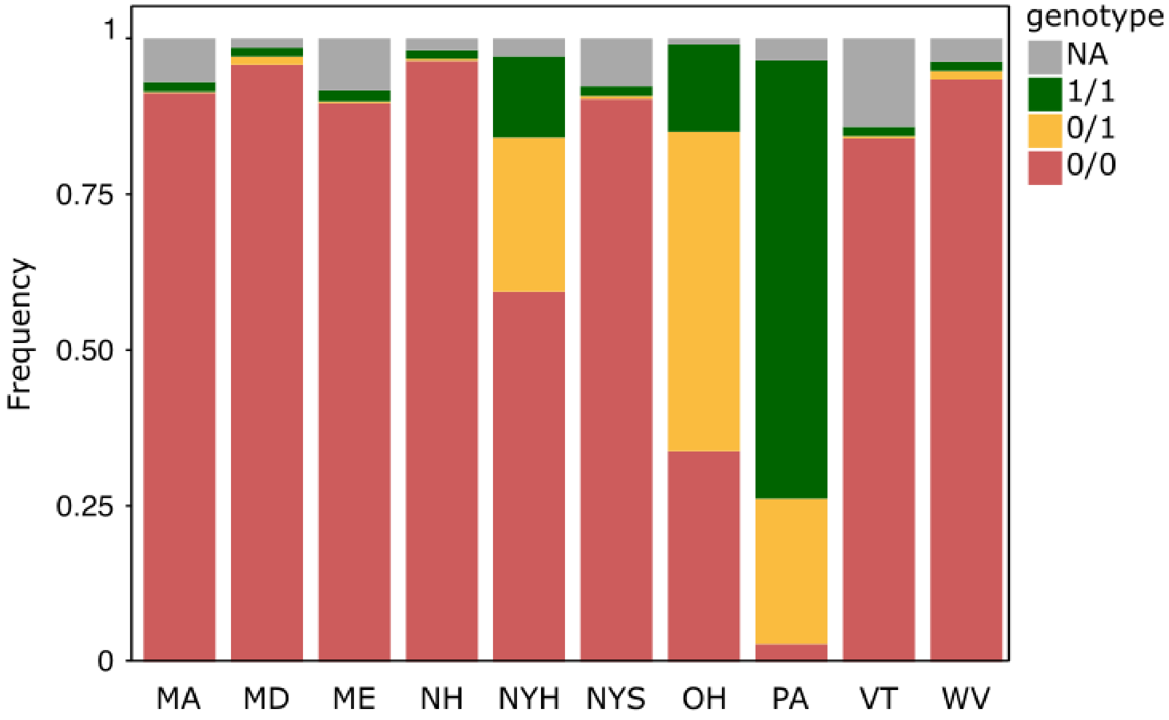


**Figure S3. Genotype Frequencies of *T. longispinosus PGRP* loci.** Boxplots shows the distribution of genotype frequencies across eleven *PGRP* genes in ten *T. longispinosus* populations. A significant effect of population origin on genotype frequency was detected (ANOVA: $F = 2.651$, $p = 0.0069$).

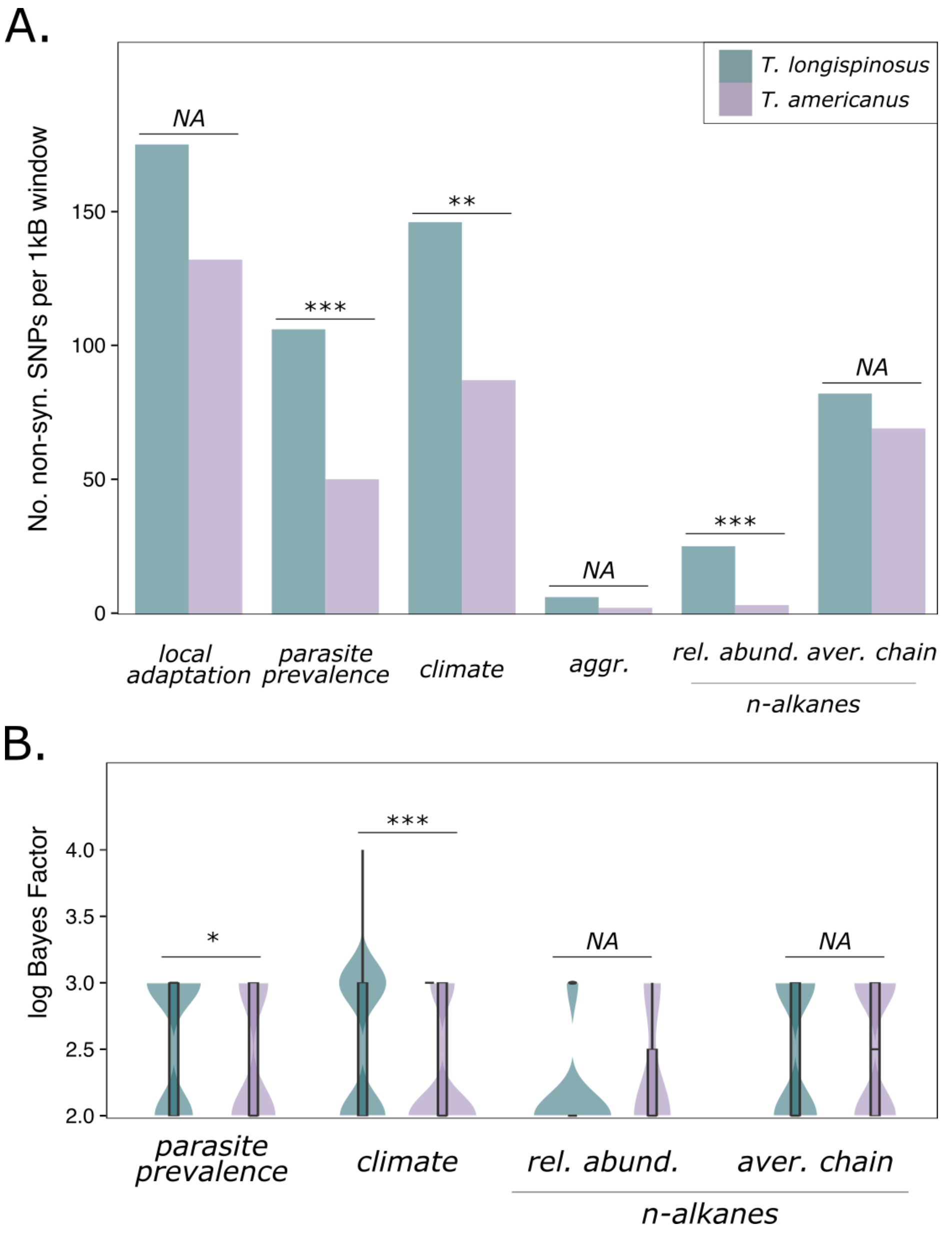


**Figure S4. (A) Number of significant non-synonymous SNPs per 1kB window per analysis and species.** The number of highly associated (BF ≥ 15) non-synonymous SNPs (ns-SNPs) within 1kB windows (i.e. unlinked) identified in each genome-wide association study. The host species *T. longispinosus* exhibited a significantly greater number of unlinked associated ns-SNPs in all analyses ($p_{adjust} \leq 0.05$) except for locally adapted loci, aggression and average *n*-alkane chain length, where both species showed a similar number of ns-SNPs. **(B) Distribution of log-transformed Bayes Factors for all ns-SNPs.** *T. longispinosus* exhibited significantly higher Bayes Factors for ns-SNPs associated with parasite prevalence ($p_{adjust} = 0.022$) and climate ($p_{adjust} = 0.0025$), but not for the relative abundance of *n*-alkanes and their average chain lengths ($p_{adjust} = 0.52$ for both).

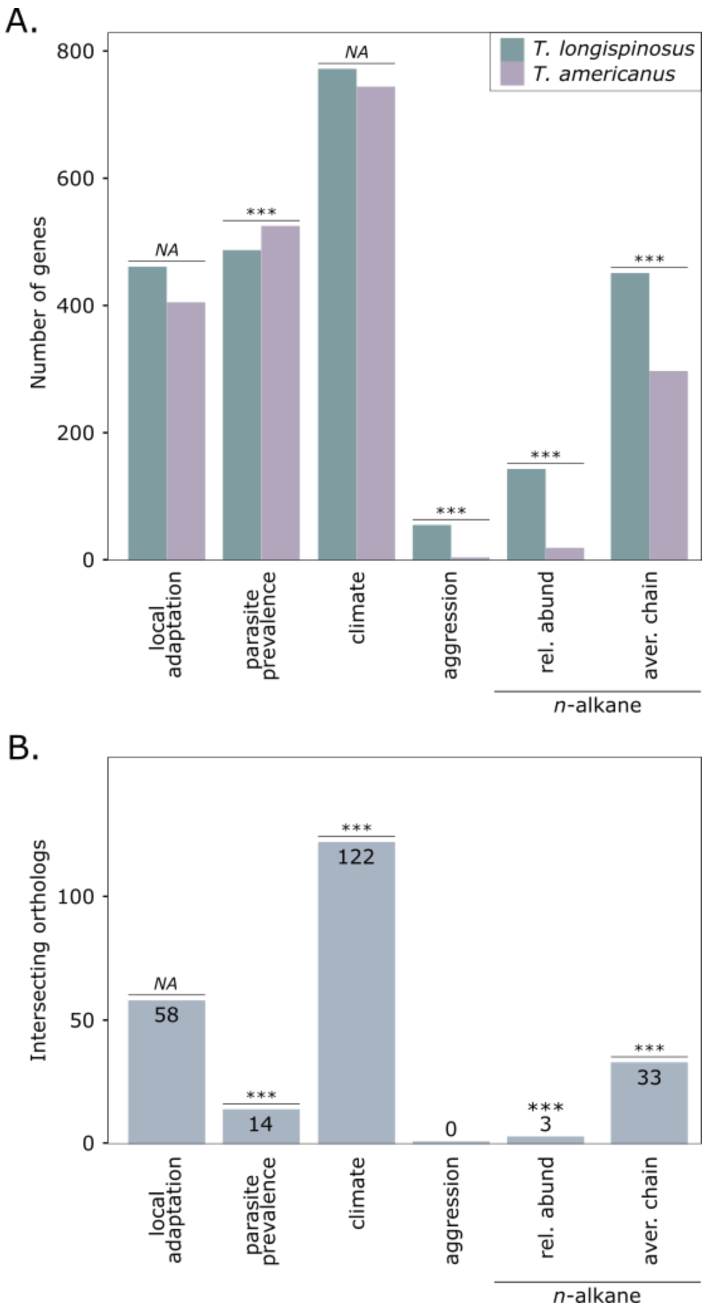


**Figure S5. (A) Number of genes containing significantly associated loci per analysis**. *T. longispinosus* exhibited significantly more candidate genes than its parasite in all analyses ($p_{adjust} < 0.001$), except for the local adaptation and climate association analysis, where both species showed similar numbers of candidate genes, and parasite prevalence associated genes, where the parasite had more genes. **(B) Orthologous candidate genes.** The number of orthologous genes identified for parasite prevalence, climate, and chemical traits was significantly greater than expected by chance ($p < 0.001$), indicating the usage of similar genes.

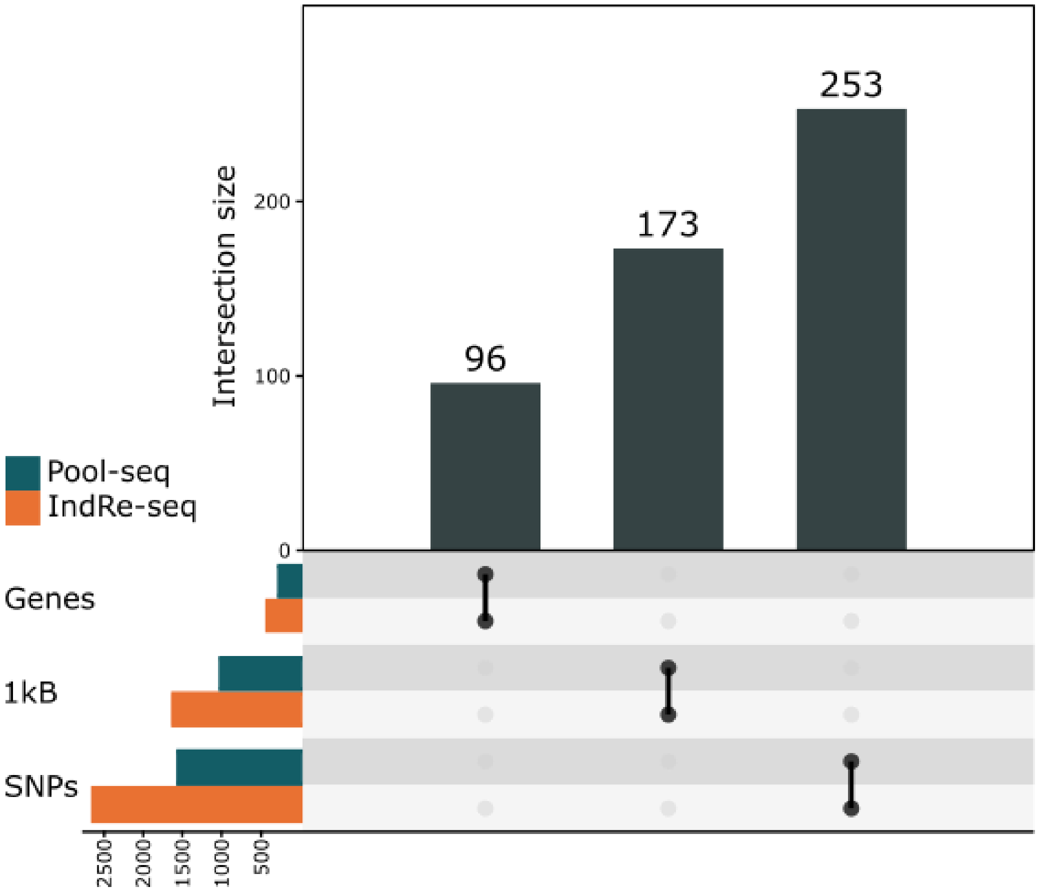


**Figure S6. Overlap of *T. longispinosus* candidate genes, genomic regions, and SNPs associated with parasite prevalence from a previous host Pool-seq study.** UpSet plot illustrating the intersections of candidate genes, unique 1kB genomic regions, and specific SNPs identified in the current host-specific parasite prevalence GWAS, as well as those previously identified using Pool-seq data (Macit et al., 2024).

#### Materials and Methods

##### *Sample Collection and Estimation of Parasite Prevalence*

The focal host species *Temnothorax longispinosus* inhabits deciduous forests in the northeastern USA and southeastern Canada (Jongepier et al., 2014). Colonies of this tiny black ant are facultatively polygynous, typically containing a few dozen workers, each measuring 2-3 mm in length. The range of this *Myrmicine* ant largely overlaps with that of its obligate social parasite, *Temnothorax americanus*, a slightly larger species with a black to brown colouration. Parasite colonies usually contain fewer than a dozen parasitic workers and fewer than 100 host workers. Colonies were collected from July to September 2021 across ten locations in the northeastern USA, with collection sites spaced 100-200 km apart (Fig. S1, Table S1). Sampling occurred near roads and tracks in state parks and on private property, with permission obtained. At each location, multiple areas within a 5 km radius were sampled to capture local diversity (Table S2). To ensure sampling of independent colonies, for both species, collected colonies had to have a minimum distance of 70 cm, and had to contain a queen. Ant colonies were transported to Mainz, Germany, maintained under standardised conditions (12:12h dark:light photoperiod at 18°C and 80% humidity), fed once per week with the Bhatkar diet (Bhatkar & Whitcomb, 1970) and given water ad libitum. Parasite prevalence serves as a measure of local parasite pressure on the host and an estimate of parasite success and was estimated as the percentage of social parasite colonies within the local *Temnothorax* community (*T. americanus* / *T. longispinosus*). Colonies of the secondary host, *T. curvispinosus*, found in MA, NYS, OH, and WV (see Table S1 for state abbreviations) were excluded. Parasitic colonies exploiting this secondary host exclusively were also excluded from calculations, and parasitic nests containing a mix of *T. longispinosus* and *T. curvispinosus* hosts were weighted based on the ratio of workers from the focal host species. Calculations were performed on long-term collection data (Table S1) (Herbers and Foitzik, 2002; Brandt and Foitzik, 2004; Foitzik et al., 2009; Jongepier et al., 2014; Kaur et al., 2019; Macit et al., 2024).

*Sample Preparation, Dissections, and DNA/RNA Extractions*

Dissections were performed in May of 2022, around eight months after collection and acclimatisation to the aforementioned standard laboratory conditions. We aimed to sample one worker from 15 independent colonies for each of the two focal species in each of the ten populations. This target was met for the host species (except for VT with 14 samples), but not for the social parasite due to low parasite prevalence in some locales. For *T. americanus,* the number of parasitised colonies sampled per population was uneven, ranging from the minimum of three samples from the NH population to 24 samples from MA (number of parasitic colonies per population: median = 19, mean = 15.4) (Suppl. S1). For the host, if possible, sampled colonies consisted of a single queen with approximately 30 workers. From each host colony, the first worker to exit the opened nest was collected to standardise sampling across the same behavioural caste. For the parasite, colonies with hosts preferably consisting only of *T. longispinosus* were chosen, although this was not always possible. There, a random parasitic worker was sampled. Ants were anesthetised on ice and quickly decapitated. The head, including antennae, was placed in a tube with 100 µl Trizol and six ceramic beads (1.4 mm diameter). The thorax, including limbs, was placed in a separate tube with six ceramic beads, and both samples were lysed for four minutes at 30 Hz using a QIAGEN Tissue Lyser II Retsch MM40. After visual inspection, additional lysis was performed for two minutes if necessary. The head/antennae samples (hereafter referred to as head samples) were then supplemented with an additional 300 µl Trizol to reach a final volume of 400 µl, and mixed to ensure complete submersion of the tissue. The fat body was carefully dissected from the abdomen and submerged in 400 µl Trizol. DNA samples were stored at -20°C afterwards, while RNA samples were incubated in Trizol at room temperature for 30 minutes before being stored at -80°C until extraction. DNA was isolated using the DNeasy Blood & Tissue Kit (Qiagen), following the insect protocol. RNA was extracted using the Direct-zol RNA Microprep Kit (Zymo) using the standard protocol. Whole-genome sequencing (WGS) and RNA sequencing (RNAseq) of 2x150 bp reads were performed by Novogene using the Illumina NovaSeq 6000 platform.

*Whole-Genome Analysis*

Sequencing data were trimmed using *Trimmomatic* v.0.39 (Bolger et al., 2014) with a 10bp head crop, and quality-checked with *FastQC* v.0.11.9 (Andrews et al., 2015). Trimmed reads were mapped using *BWA mem* v.0.7.17 (Li & Durbin, 2009) with a minimum seed length (k) of 30 and utilising unpublished reference genomes of *T. longispinosus* and *T.* americanus (Boomsma et al., 2017). This resulted in mean mapping rates of 94.4% (±2.9%) and 95.3% (±2.5%) for the two species, respectively. We used *Picard* v.2.20.8 (Broad Institute, 2018) *markDuplicates* to mark and remove duplicate sequences. We then sorted and converted to bam using *SAMtools* v.1.10 (Li & Durbin, 2009) and removed mappings with low quality using *samtools view* (-q 20 -f 0x0002 -F 0x0004 -F 0x0008). *Samtools flagstats* and *depth*, and *QualiMap* v.2.2.1 (Okonechnikov et al., 2016) were used to determine mapping rate and average coverage (see Suppl. 1). Individual BAM files were merged using *BCFtools* v1.16 (Li, 2011) with *mpileup* (--min-MQ 20 --min-BQ 13 -C50), and variant sites were called using the call function. Indels were removed, and variant sites with a depth of 5 or lower in each sample were masked. To filter out over- and underrepresented variants, the lower and upper 5% of global depth for each species were calculated and applied (925 ≥ DP ≤ 2725 for *T. longispinosus*, and 1100 ≥ DP ≤ 3114 for *T. americanus*). Finally, we excluded variants present in less than 90% of all samples (F_MISSING < 0.1) and those with a minor allele frequency below 0.05 (MAF < 0.05). With these parameters, we identified a similar number of SNPs in both species (1,677,757 SNPs for the host and 1,604,099 SNPs for the parasite). Hard-filtered vcf-files were then converted to a MAP and PED file using *VCFtools* v.0.1.17 (Danecek et al., 2011) and further converted to binary files using *PLINK* v.1.90b6.13 (Purcell et al., 2007). For population structure analyses, filtered vcf-files were further thinned by a factor of 5kB using *vcftools –thin*, resulting in around 50k SNPs per species. Since linkage disequilibrium in *T. longispinosus* is around 1kB (Macit et al., 2024), with similar linkage assumed for *T. americanus*, this window size prevented the thinned dataset from containing linked loci. All subsequent analyses were conducted on unthinned SNP data.

*Population Structure Analysis*

We performed population structure analyses using the *findstructure()* function in *sambaR* (De Jong et al., 2021) on the thinned dataset. This, among other plots, generated PCAs of SNP data by calling the *snprelate()* function (Zheng et al., 2017), and generated admixture plots that are wrapped in the *findstructure()* function. We calculated cross-entropy using the *LEAce()* function (Frichot et al., 2014) and accounted for randomness in cross-entropy calculations by running the function 50x to confidently determine the lowest k. We identified several outlier genotypes in both species and applied specific criteria to determine whether these samples should be excluded. Samples that were consistently assigned to a unique genotype in the admixture plots, from the optimal k up to k = 10, and exhibited virtually no admixture with other genotypes, were excluded from further analyses. This resulted in the removal of five samples from each species (Suppl. S1, 'sample_info'), originating from different areas, which may represent falsely identified species. Additionally, divergent genotypes were identified in both species from the Ohio population. In *T. longispinosus*, all 15 samples from Ohio were genetically distinct and thus excluded from further analyses, since their difference from all other populations might overshadow any other population-specific patterns in the GWASs. In *T. americanus*, two genetically distinct clusters were found in Ohio, referred to as parasite OH and parasite OH2 (Fig. S7B). The parasite OH cluster consisted of nine colonies with exclusively or predominantly *T. longispinosus* hosts, which were included in subsequent analyses, since samples from other populations (i.e. PA) showed admixture with this genotype. However, the more genetically distinct cluster, parasite OH2, consisted of ten colonies with exclusively or predominantly *T. curvispinosus* host workers, and was excluded due to the same reason as the exclusion of host OH. The significant genetic differences between these two clusters, despite sympatry, might suggest host specialisation or a cryptic species. Further investigations are currently underway in collaboration with C. Rabeling and M. Prebus. However, we will present population structure analysis, including this outlier OH population, further below, but not in the main text. The lowest cross-entropy was observed at k = 2 for the host when the host OH population was excluded (Fig. S2A), and at k = 3 when it was included (Fig. S7C). Similarly, for the parasite, the lowest cross-entropy was at k = 4 when the parasite OH2 was excluded (Fig. S2A), and k = 5 when included (Fig. S7C). Based on the admixture plots and the k values corresponding to the lowest cross-entropy, samples for both host and parasite were assigned to their majority genotype group as a categorical variable to determine if this grouping is associated with environmental traits. Pairwise W&C $F_{ST}$ values (Weir & Cockerham, 1984) were calculated using *sambaR*.

*Identifying outlier SNPs indicative of local adaptation*

We performed *OutFLANK* (Whitlock & Lotterhos, 2015) within the *selectionanalyses()* function in *sambaR* to identify outlier SNPs setting the do_meta flag to TRUE. This setting grouped samples according to their assigned sampled population, which allowed us to identify outlier

SNPs ($p \leq 0.05$) as a result of local adaptation/differentiation among populations. Outlier SNPs identified using *OutFLANK* will henceforth be referred to as a SNP related to local adaptation.

*Genome-wide Associations to Environment and Phenotype*

We conducted a genome-wide association study (GWAS) using *BayPass* v2.2 (Gautier, 2015) in its standard covariate mode to identify associations between single-nucleotide polymorphisms (SNPs) and two population-level environmental traits — climate and parasite prevalence — as well as two colony-level phenotypic traits — raid-related behaviours and the chemical profile. These analyses will be referred to as environmental, behavioural, and chemical GWAS. The exact values for all covariates used to create the environmental file (efile) are provided in Suppl. S2. Details of covariates for each GWAS will be explained below. Allele counts for all SNPs in all samples were extracted using *bcftools query*, specifically extracting the AD field for each sample to generate a genotype file for *BayPass*. Using allele count data rather than genotype data provides stronger association signals (González-Silos et al., 2022). A Bayes Factor (BF) of ≥ 15 was chosen as the significance threshold. Significant SNPs were categorised as within genes, in exons, in introns, 2kB up- or downstream of genes, and outside of genes using an in-house script, as well as *CROXA/tbg-tools* (Schönnenbeck et al., 2021). Special emphasis was placed on candidate genes with a high number of SNPs, specifically non-synonymous ones, and those with exceptionally high Bayes Factors (i.e., BF > 40).

*Environmental GWAS – Climate*

For climate, we used the population eigenvalues of Principal Component 1 from an analysis of climate-based data from the CHELSA bioclim database (1981-2010) (Karger et al., 2017), which included ten temperature and eight precipitation values. These PC values range from populations with warm and dry climates (negative values) to those with cold and wet climates (positive values; Fig. S1; Fig. 1A-B in Macit et al., 2024).

*Environmental GWAS – Parasite prevalence*

Parasite prevalence is defined as the percentage of parasitised colonies within the local *Temnothorax* population (*T. longispinosus* and *T. americanus* colonies), ranging from 2.3% to 15.6%. Similar to climate values, parasite prevalence is considered a population-level trait. A strong correlation was identified between parasite prevalence and climate, with warmer climates supporting a higher prevalence of parasites than colder ones. However, sufficient variability exists to distinguish between these factors in association analyses (Macit et al., 2024).

*Behavioural GWAS*

To identify the genetic basis of variations in behavioural traits relevant to both host and social parasite, we studied the response of host colonies to a sympatric invader in a preliminary study (Collin et al., in review) for which we used pivotal observed behaviours statistically linked to parasite prevalence and/or climate that were obtained around a year after collection, thus around four months after dissections of DNA and RNA samples used in this study. Around the same time, we conducted injury experiments on the host, which will be described in detail below. There, we observed differences in allogrooming behaviour in sister ants to an injured ant depending on the local climate from which the colonies originated. Both behavioural trials were conducted on ants coming from the same colonies as the ants' genotyped in this study, and were thus colony-level traits. In total, investigated behaviours included:

- Host aggression (Jongepier et al., 2014; Kleeberg et al., 2015) and brood-carrying. Only in response to parasitic invaders, host workers were noted to pick up brood and try to flee (Jongepier et al., 2015).
- Parasite aggression and passivity. Some parasites exhibited low aggression and submissive, passive behaviour (Collin et al., in review).

- Allogrooming of injured host workers. During raids, host workers are often injured or even lose limbs (Foitzik et al., 2001). The response of nestmates by grooming injured host workers covaries with climate (see below) and was thus investigated on a probable genetic basis of this behaviour.

For all behavioural GWASs, we used the total number of each behaviour performed by a colony during the trials as a covariate in the GWAS, which was normalised across the number of behaviour scans performed (Collin et al., in review).

*Chemical GWAS*

The interaction between social parasites and their hosts is closely linked to cuticular hydrocarbons, which play a crucial role in recognition within this system (Collin et al., in review). To investigate these traits and their genetic basis, cuticular hydrocarbons were extracted from two workers of a host colony and one from a social parasite colony, also used in this study, with their chemical profile being analysed by using gas chromatography-mass spectrometry (GCMS). Colony-level association analyses were performed on the following chemical information:

- *Relative abundance of recognition cues:* For both species, some methylated hydrocarbons were found to be important for eliciting aggression, including 23 CHCs for the host and 20 for the parasite (Jongepier & Foitzik, 2016; Collin et al., in review).
- *Relative abundance of (linear) n-alkanes.* Linear *n*-alkanes are generally not used in recognition but serve to protect against desiccation. Social parasites exhibit a profile rich in *n*-alkanes (Kleeberg et al., 2017), which might help them to avoid host detection (Kaur et al., 2019).
- *Average chain length of n-alkanes.* Longer-chained *n*-alkanes have better desiccation protection capabilities, as longer chains form longer and tighter layers. Since we

identified population- and species-specific differences in their length (Fig. S10), a genomic basis for this chemical trait could be assumed.

The number of samples included in each GWAS varied slightly among the environmental, behavioural, and chemical GWASs, as well as between species, due to the inability to consistently obtain both genotype and phenotype data for all samples (summarised in Table S3).

*RNA analysis*

Trimming and quality control of RNA-seq data were performed as described above for WGS data. Reads were mapped using *HISAT2* v.2.1.0 (Kim et al., 2015) to the aforementioned reference genome. Mean mapping rates for transcriptome data in *T. longispinosus* were 64.2% (±33.4%) for fat body samples and 80.1% (±26.3%) for head samples, and in *T. americanus* 64.9% (±31.5%) for fat body samples and 80.9% (±22.7%) for head samples. Using the *htseq-count* function from *HTSeq* v.2.0.2 (Putri et al., 2022), we created transcript read count tables used as input for *DESeq2* v.1.42.0 (Love et al., 2014) in *R* v.4.3.2 (R Core Team, 2021). Transcripts with less than ten counts in at least one-third of all samples were removed. Due to high variances in all four RNAseq datasets, we filtered for outliers using *pca.outlier()* function from the *mt* package v.2.0.1.19 (Lin, 2021), which identifies outliers based on Mahalanobis distance of PC1 and PC2 values. Outlier detection was performed twice for all four read count tables, and outlier samples were removed from the original read count table (see Suppl. S1). The read count tables were then used to conduct a PCA. Differentially expressed genes (DEGs) were tested using *DESeq2* with variance stabilised data, with each parasite prevalence (results presented in the main manuscript), climate PC1 eigenvalues and aggression values (results presented here) used as continuous variables in both tissues and both species. We identified DEGs that contain loci in putative promoter and intron regions associated with parasite prevalence, as these may impact expression patterns (Cooper, 2010).

*Functional and Comparative Analyses*

To further characterise candidate genes (i.e. those containing significant SNPs in any of the analyses), we obtained functional annotations for all genes in the reference genomes using their peptide sequences. First, we ran *InterPro* v.5.61.93 (Paysan-Lafosse et al., 2023) to retrieve Gene Ontology (GO) information, which was used to perform functional enrichment analyses using *topGO* v.2.54.0 (Alexa & Rahnenfuhrer, 2017). Next, we ran a *BlastP* v.2.13.0 (Altschul et al., 1990) search against the non-redundant invertebrate database (retrieved on NBCI on Jan 2022) and also a proteome database consisting of *Drosophila melanogaster* and *Apis mellifera* (retrieved on UniProt on Jan 2024; Proteome IDs: UP000000803 and UP000005203) (The UniProt Consortium, 2023). Any gene names provided in this study for specific candidate genes will always refer to the best BLAST hit generated in *A. mellifera*, unless stated otherwise. We further retrieved the UniProt entry ID, associated GO terms, function text and associated publications via the UniProtKB entry for hits in *A. mellifera*. Orthologs between both species were identified using *OrthoFinder* v.2.5.4 (Emms & Kelly, 2015), and their GO functions were summarised using *REVIGO* (Supek et al., 2011). To test whether the number of candidate genes in each GWAS and differentially expressed genes differed between species, we used the built-in chi-square function in *R*, considering the total number of genes in both reference genomes (*T. longispinosus* = 16,064, and *T. americanus* = 14,128). Similarly, to test whether the number of non-synonymous SNPs differed between species, we used the chi-square test, considering the total length of exon regions in both reference genomes (*T. longispinosus* = 21,924,724 and *T. americanus* = 20,133,265). For non-synonymous SNPs, we tested for significant differences in log-transformed Bayes Factors between species using a one-way ANOVA followed by a Tukey HSD post-hoc test to identify pairwise differences. To test if the number of overlapping orthologous candidate genes in each GWAS and differentially expressed genes differed between species, we used the *hypergamous()* function from the Python-based *SciPy* software (Virtanen et al., 2020). For all statistical tests, we chose a significance threshold at FDR-corrected $p \leq 0.05$ and Bayes Factors (BF) $\geq 15$.

*Demographic History Analysis*

We employed a pairwise sequentially Markovian coalescent model (PSMC; Liu & Hansen, 2017) to detect changes in the effective population size ($N_e$) over a broad temporal scale. We chose two samples per population per species with the lowest level of admixture (excluding host OH and parasite OH2) (Suppl. S1). We created mpileup files per species and called variances using *bcftools*. Consensus files were generated using *vcfutils.pl vcf2fq* (-d 10 -D 100) and converted to the appropriate data type using *fq2psmcfa*. The PSMC analysis was performed across 64 interval times (-p 6+25*2+4+4) and the upper limit of the TMRCA (-t) set to 10. Plots were generated using *psmc_plot.pl* with generation time set to 10 years (-g 10) as was previously described in the host species (Kaur et al., 2019) and mutation rate set to the one found in *A. mellifera* (-μ 3.4e-9; Liu & Hansen, 2017; Yang et al., 2015) as was similarly done in *Solenopsis* ants (Cohen & Privman, 2019).

#### Additional Results and Discussion

*Population Structure*

A principal component analysis (PCA) of the thinned SNP dataset for the host revealed no clear clustering by population, although samples from Massachusetts (MA) were distributed toward the far negative side of the PC1 axis, and those from Maine (ME) toward the far positive side (Fig. 1A). Admixture analysis indicated that the lowest cross-entropy occurred at $k = 2$ ancestral populations (Fig. S2A). Populations from MA, New York Huyck (NYH), New York South (NYS), and West Virginia (WV) were predominantly characterised by the blue genotype (genotype 1), whereas ME was primarily associated with the green genotype (genotype 2; Fig. 1B). Each sample was assigned to its majority genotype, with these assignments being significantly influenced by local parasite prevalence (ANOVA: $F = 42.72$, $p < 0.0001$), climate values ($F = 30.75$, $p < 0.0001$), and their interaction ($F = 6.57$, $p = 0.012$; Fig. S2B). For the parasite, the PCA of thinned SNP data showed clearer population clustering, in particular in samples collected in OH, Maryland (MD), WV, and Pennsylvania (PA; Fig. 1A). Admixture analysis revealed the lowest cross-entropy at $k = 4$ (Fig. S2A), with distinct genotype patterns observed in MA, OH, Vermont (VT), and WV (Fig. 1B). Each sample was assigned to its majority genotype (blue, green, red and orange, Fig. 1B). These assignments were significantly affected by local parasite prevalence (ANOVA: $F = 46.84$, $p < 0.001$), climate values ($F = 9.01$, $p = 0.0032$), and their interaction ($F = 29.43$, $p < 0.0001$; Fig. S2B).

*Population Structure including outlier Ohio populations*

Our population structure analysis of the primary host, *T. longispinosus*, revealed stark differences in all host samples from Ohio (referred to as 'host OH' henceforth) compared to those from all other sites. Similarly, in the parasite, samples from OH clustered into two distinct groups (referred to as 'parasite OH' and 'parasite OH2' henceforth), seen as two distinct clusters in a PCA of thinned SNP data (Fig. S7A). The same grouping was also evident in admixture plots, where parasite OH2 showed a distinct genotype with no admixture in any other samples or populations (Fig. S7B). The lowest cross-entropy in the dataset, with and without host OH and parasite OH/OH2, differed by 1, indicating that host OH and parasite OH2 consistently formed their own ancestral groups (Fig. S7C). Pairwise $F_{ST}$ values in host and parasite were slightly higher when host OH and parasite OH2 were included (Fig. S7D). Isolation-by-distance patterns were slightly stronger in the host ($p = 0.089$), and in the parasite ($r = 0.38$, $p = 0.0016$; Fig. S7E). Our metadata revealed that *T. americanus* samples with the parasite OH genotype (blue in Fig. S7B, $k = 5$) and the parasite OH2 genotype (purple) differed in the host species they enslaved: Parasitic colonies with majority parasite OH genotype almost exclusively enslaved *T. longispinosus* hosts, and those with the parasite OH2 genotype enslaved predominantly *T. curvispinosus* hosts or a mix of *T. longispinosus* and *T. curvispinosus* hosts. For the host, we were unable to identify a driver for the unique host OH genotype based on our metadata. Further research on possible host-driven speciation, or identification of a cryptic parasite species, should increase sample sizes of host and parasite, noting down the host species in parasitised colonies, and expanding sampling sites to the US states of Illinois and Michigan, which contain *T. curvispinosus* in high densities.

Tajima's D values were consistently low and slightly positive across populations in both *T. longispinosus* and *T. americanus*, reflecting a mild excess of intermediate-frequency alleles (Fig. S8). This pattern arises from nucleotide diversity ($\pi$) exceeding Watterson's theta ($\theta$) in all populations, suggesting stable population sizes with no recent bottlenecks or expansions in most populations. The moderate and tightly clustered $\pi$ and $\theta$ values indicate relatively high genetic diversity and demographic uniformity across regions in both species. The similarity between host and parasite implies shared evolutionary dynamics, possibly due to common environmental pressures or interdependent dispersal histories, though the low magnitude of Tajima's D points to weak or background-level selective forces rather than strong directional or balancing selection.

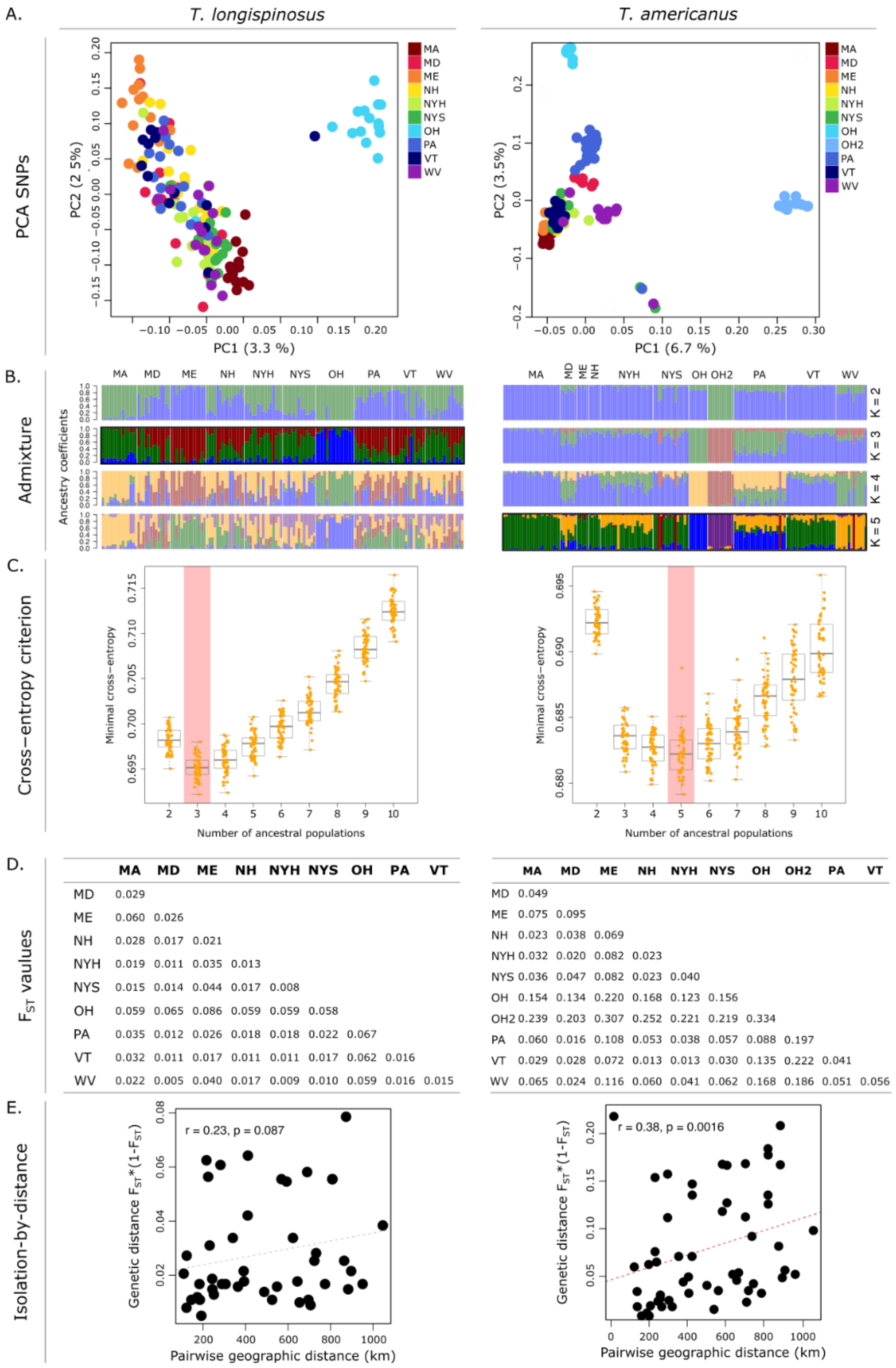


**Figure S7. Population structure analysis including all Ohio samples.** (A) Principal Component Analysis of SNP data, based on a thinned dataset containing ~50k SNPs per species. (B) Admixture plot depicting the various genotypes identified and their distribution/proportion across samples in populations. Highlighted is the optimal k. (C) The determination of the optimal number of ancestral populations (k) based on the cross-entropy criterion, revealing k = 3 for *T. longispinosus* and k = 5 for *T. americanus* using LEA() ran 50x. (D) Pairwise $F_{ST}$ values. (E) Isolation-by-distance.

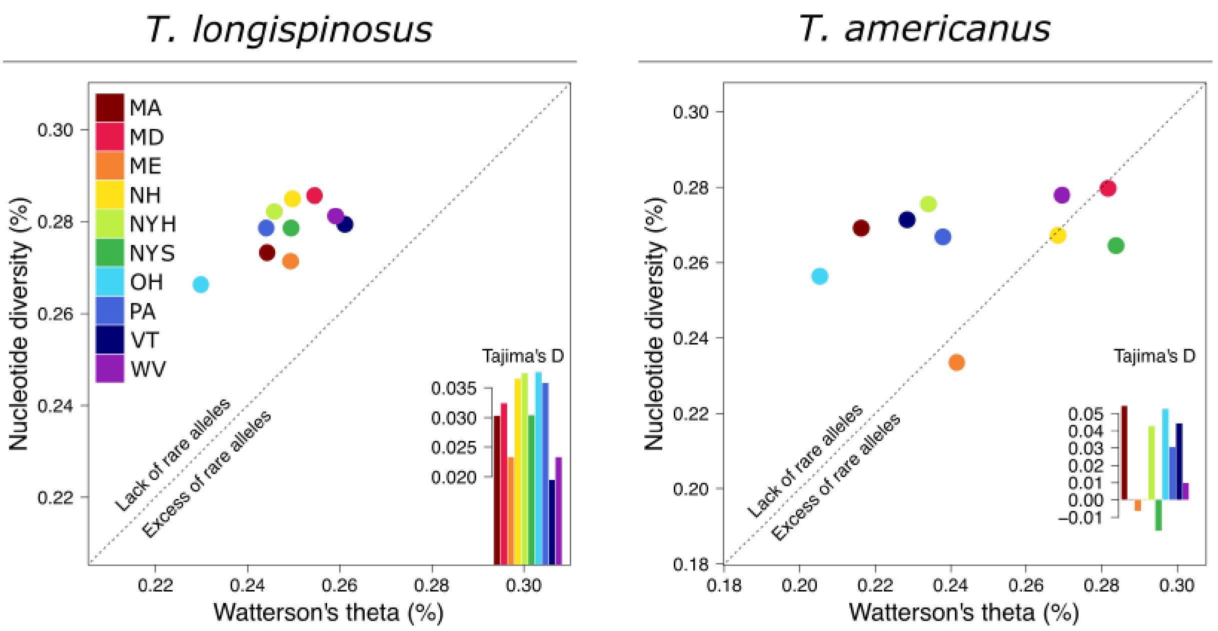


**Figure S8. Comparative SNP frequency spectrum**. Scatterplots show the relationship between nucleotide diversity (π) and Watterson's theta (θ) for each sampled population in both species. The dashed line represents π = θ, along which Tajima's D = 0, with points above the line indicating a relative deficit of rare alleles (positive Tajima's D), while those below indicate an excess of rare alleles (negative Tajima's D). Insets display population-specific Tajima's D values, highlighting consistently positive values in *T. longispinosus* (left) and more variable patterns in *T. americanus* (right), suggesting differences in demographic history or selective pressures between host and parasite, but are likely additionally the result of uneven and small sampling sizes in the parasite.

*Demographic History Analysis*

Our PSMC analysis yielded counterintuitive results, with the $N_e$ of the parasitic ant species being higher than that of its host: *T. americanus*, as a dulotic parasite, is usually rare, resulting in lower population sizes, as reflected in its classification as a vulnerable species on the IUCN Red List (IUCN, 2022). While the absolute $N_e$ values may underestimate true population sizes, particularly in haplodiploid social insects where $N_e$ is calculated differently than by PSMC

(Hedrick & Parker, 1997; Cohen & Privman, 2019), the observed disparity warrants further discussion. A potential explanation for this discrepancy is the substantially higher worker reproduction rate in the parasite. Host worker reproduction is restricted to queenless colonies (Bourke, 1988). In contrast, it is a regular and significant component of the parasite's reproductive strategy, with studies reporting up to 70% of parasitic males being worker-reared in *T. americanus* (Buschinger & Alloway, 1977; Foitzik et al., 2001). Given the parasite's prevalence (2-15%), which suggests a comparatively smaller population size, the high rate of worker reproduction could contribute to a higher-than-expected $N_e$. Furthermore, while the parasite exhibits clear population structuring, potentially leading to an overestimation of $N_e$, the host appears closer to panmixia, which could lead to an underestimation of $N_e$ (Hilgers et al., 2024).

Both species exhibit a peak in $N_e$ around 2 million years ago, but their trajectories diverge, with the parasite's population sizes dropping seemingly more rapidly than those of the host, both reaching similar values around 100,000 years ago (Fig. S9). The Pleistocene ice age in the Appalachian Mountains likely prompted southward migrations, with certain areas within our sampling range (PA, MD, WV and partially OH) serving as ice-free refugia (Bloom, 2018; Earth@Home, 2024). The subsequent glacial retreat around 11,000 years ago likely facilitated recolonisation and significantly influenced contemporary population structures. These ice-free areas also have high parasite prevalence, which may suggest the parasite inhabits these regions for longer compared to northern populations.

Finally, the host and parasite differ in major population traits, with low population structure in the host and strong structuring in the parasite. This may result in problems comparing the results of both species. We acknowledge those and other limitations associated with PSMC analyses, including sensitivity to parameter choice and potential inaccuracies when using scaffold-based genomes as was done here, particularly for recent time scales (Li & Durbin, 2011). Therefore, the PSMC results presented here should be interpreted cautiously.

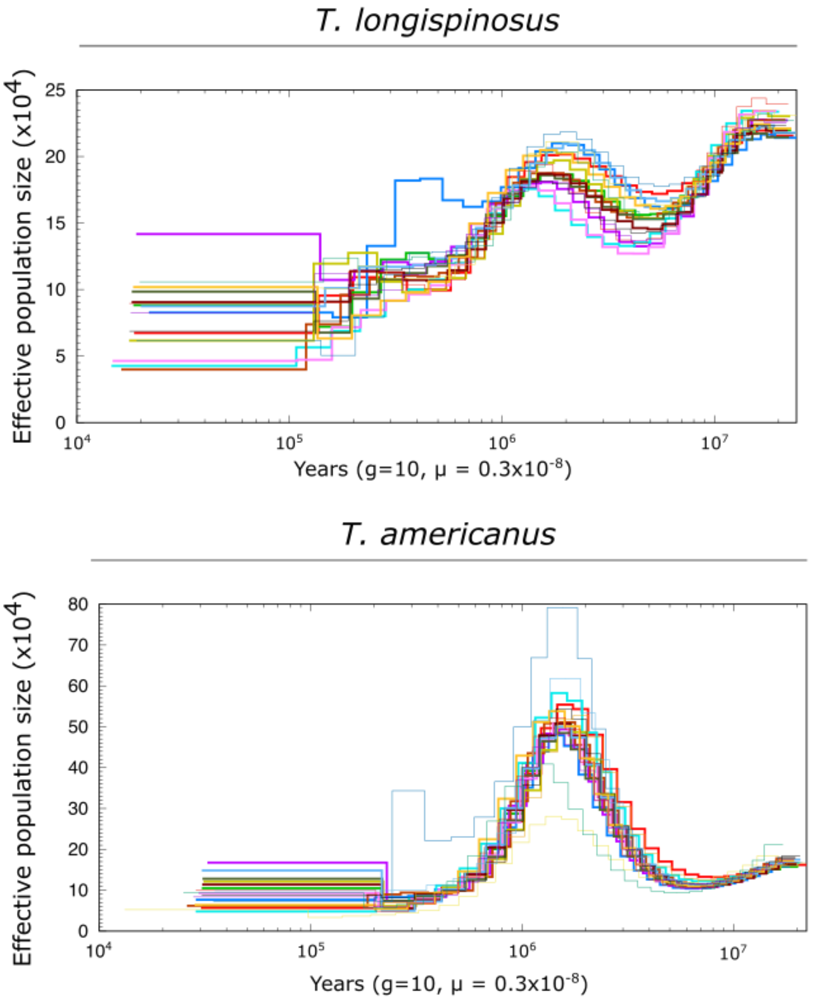


**Figure S9. Demographic history analysis.** Parameters used were -p 6+25*2+4+4, setting the upper limit of the TMRCA (-t) to 10, generation time to 10 years (-g 10) and mutation rate to 3.4e-9 (-μ). Effective population size ($N_e$) over time for host and parasite shows peaks around 2 million years ago, followed by a steeper decline in parasite $N_e$.

*Chemical Characteristics*

The chemical composition of the cuticular hydrocarbons in social insects serves a dual function, protecting against desiccation and facilitating communication (Sprenger et al., 2019). The length of linear *n*-alkanes can influence their protection against desiccation properties, with longer linear *n*-alkanes creating tight layers with low viscosity. Conversely, shorter chains allow better fluidity through the cuticle of insects (Menzel et al., 2019). The average chain lengths in both species appear to be similar, with small standard deviations (host: average 27.74, std: 0.16; parasite: 27.28, std: 0.12). We performed a one-tailed ANOVA test using *aov()* function in *R* and confirmed that variances in average chain lengths of linear *n*-alkanes are significantly different between populations in the host ($Df = 8$, $F = 3.504$, $p = 0.0012$) and in the parasite ($Df = 9$, $F = 3.668$, $p = 0.00050$), and highly differed between species, with the parasite showing shorter chain lengths than the host (Mann-Whitney U test: $z = 13.62$, $p < 0.00001$; Fig. S10), which has been reported before in this and other dulotic ant species, and discussed to reduce host detection (Kleeberg et al., 2017). Based on these observed differences, we felt justified in investigating the putative genomic basis of these significant differences in chain lengths and thus performed a GWAS, for which we report the results in the main document.

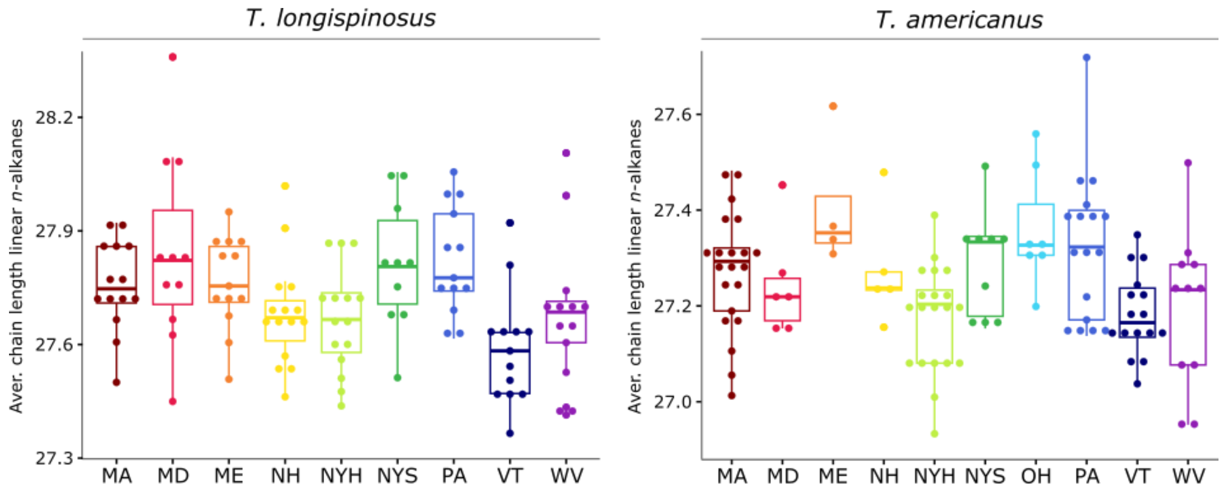


**Figure S10. Linear *n*-alkane chain length variations.** Boxplots showing the distribution of average chain lengths of linear *n*-alkanes in host and parasite populations. Significant differences in chain length were observed within both host (ANOVA, $p = 0.0012$) and parasite ($p = 0.0050$) populations, as well as between species (Mann-Whitney U test, $z = 13.62$, $p < 0.00001$).

*GWAS of species-specific behaviours*

In Collin et al. (in review), host and parasite ants exhibited specific behaviours during experimental raids that appear adaptive to local environmental pressures. The host ants exhibited aggressive behaviour toward parasites, akin to 'fight', and brood-carrying behaviour, which aligns with 'flight' or protective responses. Parasites, in contrast, showed aggression and passive 'freeze' behaviours, with inactivity potentially serving as a survival tactic. The primary objective of this study was to explore the genetic underpinnings of co-evolutionary traits in these interactions, with a particular focus on aggression, given the availability of reciprocal data between hosts and parasites. However, given the observed associations between other species-specific behaviours and parasite prevalence, a GWAS was conducted on these behaviours and is reported in the following.

*Host – brood-carrying*

We identified six genes containing significant loci associated with brood-carrying behaviour in the hosts, including one non-synonymous SNP (ns-SNP) in a *FAS* gene (Suppl. S2). Two other candidate genes were *IDE* and *radish*, which has roles in olfactory long-term memory (Folkers et al., 2006). In the host aggression GWAS, *IDE* was suggested as influential in age-dependent aggression, given its role in other insulin-related genes in the shift from nursing to foraging (Chen et al., 2023). Notably, foragers, rather than nurses, typically engage in parasite defence (Koenig & Moreau, 2024). Since brood-carrying is primarily a nursing behaviour, the findings imply a complementary role for *IDE* in age-specific host responses to parasitic intruders. Additionally, *radish* and *FAS* may facilitate recognising and memorising parasite-associated chemical cues, initiating brood-carrying to protect vulnerable larvae and pupae.

*Parasite – passive*

We identified five genes containing significant loci associated with passive 'freeze' behaviours in the parasite, although none were non-synonymous (Suppl. S2). Noteworthy genes include an *odorant receptor* gene, and those involved in transcription and gene expression (*histone demethylase UTY*, *polyglutamine-repeat protein pqn-41*). The number of candidate genes relating to gene expression and transcription might suggest a stronger emphasis on gene expression patterns for this specific behaviour. Earlier research has shown that parasites exhibit shifts in gene expression when transitioning from a resting to a raiding state (Alleman et al., 2018), underscoring the selective pressure on gene regulatory functions. Further and more generally, passive or inactive behaviour within its parasitic nest was previously deemed to be indicative of 'behavioural water-saving strategies' in support of their strategy of chemical insignificance suggested by Lorenzi (2021), and might link passiveness to a default behaviour.

*Differentially Expressed Genes associated with Climate and Orthologous Gene Expression*

We performed *DESeq2* on fat body and head samples for both species using climate PC1 eigenvalues as a continuous variable. For the host fat body transcriptome, we identified 525 genes to be significantly associated with climate PC1 values, which included several *FAS* genes. For the parasite's fat body transcriptome, we identified significantly more genes (884; gene count: $\chi 2 = 95.13$, $p_{adjust} < 0.001$), including candidate genes such as *chitinase domain-containing protein 1 (CHID1)* and a *fatty acyl-CoA reductase (FAR*; Fig. S11, Suppl. S1). The host fat body transcriptome showed 18 enriched functions, with many relating to mitochondrial functions (i.e., 'mitochondrial respiratory chain complex I assembly', 'mitochondrial electron transport') and meta- and catabolic processes (i.e., 'proteolysis involved in protein catabolic process', 'galactose metabolic process'). For the parasite fat body, we identified 20 enriched functions, with most of them being involved in either gene expression itself ('translation', 'mRNA transport', 'protein transport', 'protein folding') or meta- and catabolic processes (i.e., 'cellular metabolic process', 'catabolic process'; Suppl. S1). For the host's head transcriptome, we identified 49 genes significantly associated with climate PC1 values, most notably an *odorant binding protein* (*OBP*) gene and a *phosphodiesterase* (*PDE*). The parasite's head transcriptome showed a similar number of genes (40; gene count: $\chi 2 = 0.72$, $p_{adjust} = 0.44$), and similarly included an *OBP* gene (Fig. S11, Suppl. S1). The host head transcriptome showed nine enriched functions, with half of them being involved in metabolic processes (i.e., 'glucose metabolic process', 'lipid metabolic process'). The parasite's head transcriptome showed 10 enriched functions, with the most enriched ones relating to gene expression processes (i.e., 'translation', 'regulation of mRNA processing'; Suppl. S1).

Sympatric host–parasite populations are exposed to the same environmental conditions (parasite prevalence and climate). We therefore identified orthologous genes that are differentially expressed in response to these environmental conditions in both species, indicating a similar utilisation of gene expression pathways. For orthologous expressed genes associated with parasite prevalence in both species' fat bodies, we identified 21 differentially expressed orthologous genes associated with parasite prevalence, which was more than expected by chance (hypergeometric test: $p < 0.001$, Jaccard index = 0.034; Suppl. S1). These included genes with functions in oxidative stress and neuronal activity such as c*uticular protein 14 precursor*, s*uperoxide dismutase*, *proteasome subunit alpha type*, and *clavesin-2*. We used the online tool *REVIGO* to identify commonalities among the associated Gene Ontology (GO) terms of orthologous expressed genes. We found them to be related to core cellular processes ('cellular component organization') and gene regulatory functions (i.e., 'regulation of transcription by RNA polymerase II'). For orthologous expressed genes associated with climate for both species' fat bodies, we identified 77 genes, which was more than expected by chance (Jaccard index = 0.061, $p < 0.001$). Among those were genes with functions in fatty and lipid metabolic processes (*very long-chain-specific acyl-CoA dehydrogenase*, *long-chain-fatty-acid-CoA ligase*), but also cuticle composition (*probable chitinase 10*; Suppl. S1). Using *REVIGO*, we identified commonalities in associated GO terms, of which the vast majority were related to gene regulatory functions (i.e., 'regulation of DNA-templated transcription', 'mRNA processing'), and further some post-translational modifications ('protein phosphorylation', 'protein dephosphorylation'). For both environmental factors, parasite prevalence and climate, we found no associated orthologous expressed genes in head samples.

In our transcriptome analyses, using parasite prevalence as a continuous variable, we consistently observed significantly lower transcriptional activity in the parasite compared to the host (see main text). However, using climate as a factor revealed a partial reversal of this pattern, with the parasite now exhibiting increased constitutive gene expression in its fat body associated with climate than its host. We also found more orthologous expressed genes between the two species related to climate than parasite prevalence, using shared genes as pathways involved in gene regulatory functions. Investments in constitutive gene expression of the head transcriptomes further suggest similar climate-driven transcriptional adaptations in this organ across both species. Functional enrichment analyses revealed the use of similar processes and pathways across tissues and species: Metabolic and catabolic processes were enriched in the fat body transcriptomes of both the host and parasite, as well as in the host's head. Mitochondrial functions were also enriched in the host fat body, while gene expression-related functions were prominent in the parasite fat body and head transcriptomes. These enriched metabolic/catabolic, and mitochondrial/ATP-related functions may reflect a shared investment in processes that scale with temperature, potentially influencing growth and body size in both species, as is the case in other insects (Bjørge et al., 2018). The identification of *FAS* and *FAR* genes in both species may also reflect climate-adaptive chemical desiccation protection, given their role in the CHC biosynthesis. Intriguingly, *OBP* gene were identified as candidate genes in both species in their head samples: This finding may indicate an adaptive response to increased volatility of chemicals in warmer temperatures, potentially enhancing odorant perception (Biessmann et al., 2010), including, the detection of alarm pheromones (Du & Chen, 2021).

*Differentially Expressed Genes associated with Aggression*

Using aggression values obtained from Collin et al. (in review) for identifying differential gene expression, we identified 313 DEGS in the host's fat body and none in its head in association with aggression. In contrast, the parasite exhibited 2,701 DEGs in the fat body and two in the head (Suppl. S1). Given the negligible number of DEGs in head samples, subsequent analyses focused exclusively on fat body tissue. There, the parasite displayed more than eight times the number of DEGs associated with aggression compared to its host, a statistically significant difference (gene count: $\chi 2 = 3269$, $p_{adjust} < 0.0001$). Functional enrichment analysis revealed 17 enriched biological GO terms in the host and 32 in the parasite. Both species shared two enriched terms related to gene regulatory processes (‘translation’, ‘translation initiation’). In the host, additional enriched functions included further regulatory processes (‘mRNA export from nucleus’) and post-translational modifications (‘post-translational protein modification’). Nearly all enriched functions in the parasite were related to gene regulation (‘mRNA splicing via the spliceosome’, ‘ribosomal large/small subunit biogenesis’) (Suppl. S1). However, due to the low number of aggression data available, this analysis was performed on around 1/3$^{rd}$ of the available samples, and results should be interpreted cautiously.

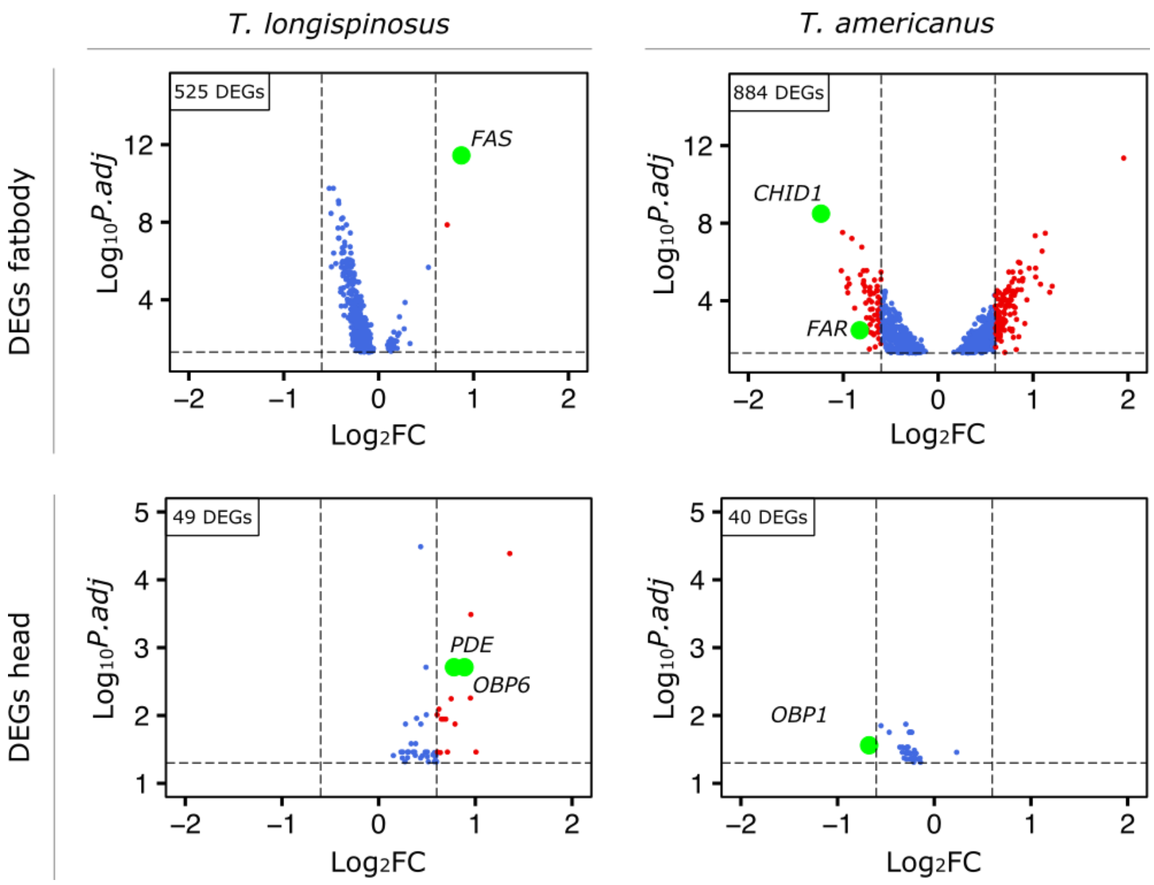


**Figure S11. Differential gene expression in *T. longispinosus* and *T. americanus* in response to climate.** Volcano plots showing differential gene expression in fat body and head tissues of *T. longispinos*us and *T. americanus* associated with climate PC1. Plotted are significantly differentially expressed genes (DEGs; $p \leq 0.05$) with thresholds drawn at $\log_2 FC = 0.7$. Selected candidate genes of interest are labelled.

### First Aid in Ants: Response of *Temnothorax* Ants to Injury in Relation to Social Parasite Prevalence and Climate

*Abstract*

Destructive raids by dulotic parasitic ants often injure defending host ants, prompting nestmates to provide wound care to reduce the risk of secondary infections. To test whether this behaviour varies with parasite pressure, we simulated raid-induced injuries in *Temnothorax longispinosus*, host of the dulotic parasite *T. americanus*. Injured ants from warmer climates received more frequent and prolonged allogrooming, suggesting a temperature-driven adaptation to potentially elevated microbial risks. In contrast, grooming decreased in populations with higher parasite prevalence, possibly reflecting a strategic shift toward replacing injured workers instead of caring for them. These results highlight the interaction between climate and parasitism in jointly shaping social care strategies, revealing context-dependent adaptations in host–parasite systems.

*Introduction*

Injuries can pose high risks by either being fatal or indirectly via subsequent infections (Hart, 2011), with further consequences on colony-level in social insects. Extreme behaviours, such as self-amputation (Emberts et al., 2017) and the amputation of injured limbs by sisters in ants (Frank et al., 2024), have evolved to reduce injury costs or infection risks. The socially parasitic ant *Temnothorax americanus* replenishes its workforce through destructive raids on host colonies, often injuring host workers. Given the varying prevalence of *T. americanus* (2-16%) across the interaction range with its host, *T. longispinosus*, we investigated whether the host's injury response via leg amputation varies with parasite pressure, but given its ecological impact on parasite prevalence, we further included climate as an explanatory factor (Macit et al., 2024; Collin et al., in review). In early 2024, we conducted injury experiments on *T. longispinosus* colonies (min = 1, max = 18, mean/median = 13 colonies per population; Fig.

S12) from our existing host–parasite collection. By creating injuries via dissection of a limb and subsequently assessing the behaviour of injured ants and their sisters, we attempted to examine the relationship between host responses to injured nestmates along a parasite prevalence and climate gradient, as well as in relation to the original host colony size. We predicted that (i) hosts in highly parasitised areas show increased injury care, potentially due to selection on immune genes in these populations (Macit et al., 2024) and (ii) warmer climates might favour increased grooming of injured nestmates to reduce infection risks of highly proliferating bacteria in warmer temperatures (Linder et al., 2008). Ant behaviour was scored using BORIS software (Friard & Gamba, 2016) based on an established ethogram (Table S7).

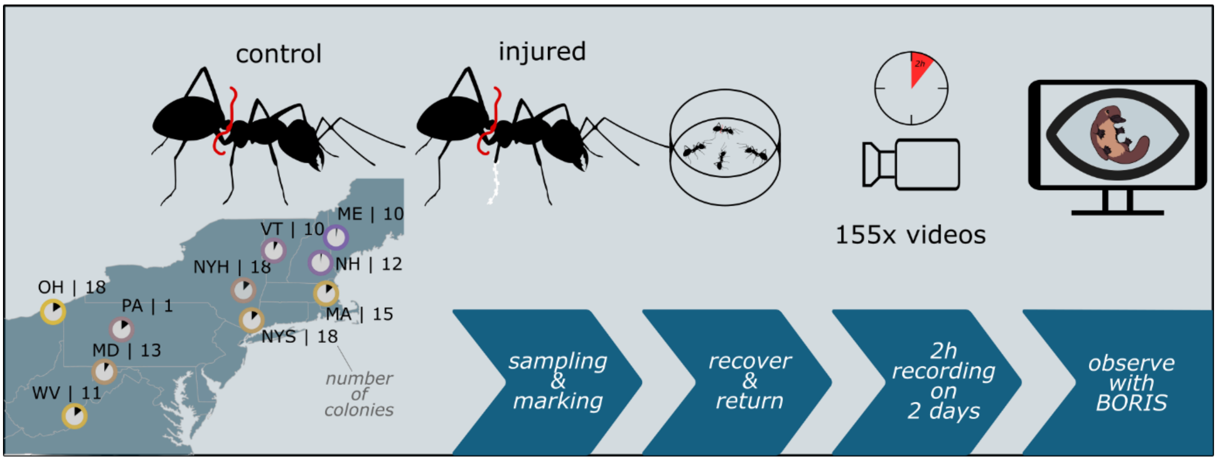


**Figure S12. Experimental design for injury assays.** Healthy host colonies from each population were selected. A worker ant was randomly selected and marked with a red wire. For the treatment groups (injured and non-control), the middle leg was quickly removed. Following a 5 minute recovery period, both control and injured ants were returned to their original colonies, and their interactions were recorded for a period of 2 hours. This observation was repeated for an additional two hours on the following day if the control or injured ant survived.

*Materials and Methods*

Colonies were collected and maintained as described in the Materials and Methods section above. In total, one to 18 independent host colonies per population were used (see map in Fig. S12). The number of queen(s), workers and larvae was counted before the experiment.

Coloured wire loops were used to mark ants to be injured, and their controls. Controls and experimental ants were briefly sedated with $CO_2$. For the experimental ants, a single middle leg was cut using sterilised micro scissors, followed by a 5 minute recovery period before reintroduction to the nest for observation. This procedure was similarly applied to the control ants with no leg amputation. Recordings lasted two hours, with each colony observed for two consecutive days if the injured ant or the control survived the first day. Video analysis was conducted using *BORIS* software (Friard & Gamba, 2016), focusing on the behaviours of injured and control ants as defined in an ethogram (Table S7). We quantified the occurrence of observed behaviours, activities, and interactions, and for a subset of these, their duration was also measured (Suppl. S3). Data analyses involved generalised linear mixed models (*glmms*) using the *glmmTMB* package (Brooks et al., 2017) in *R* v.4.2.3 (R Core Team, 2021). These included the behaviour as dependent variables, the treatment (control vs. injured), parasite prevalence, climate PC1, the interactions between the treatments and each of these ecological factors, and colony size as a random factor. Models were checked for residual fit using the *DHARMa* package (Hartig, 2016), given a distribution that seemed adapted to the type of data. Model reduction was performed based on the assessment of multicollinearity using the *performance* package (Lüdecke et al., 2019) to eliminate parameters causing collinearity, followed by stepwise automatic model reduction using the *buildmer* package (Voeten, 2019) based on the Akaike Information Criterion (AIC) to obtain the final models. The output of the resulting models was extracted using the *emmeans* package (Lenth, 2025) with the *joint_tests()*, *emmeans()*, and *emtrends()* functions to obtain the estimates.

**Table S7. Ethogram of observed behaviours.** List of observed and scored behaviours during the experiments, including abbreviations and detailed descriptions of each behaviour.

| Behaviour | Abbreviation | Description |
|---|---|---|
| *Allogrooming* | G1 | Interaction of sister ant with the injury or the control leg |
| *Selfgrooming* | G2 | Selfgrooming the injury or the control leg |
| *Antennating* | A1 | Antennating the injury or the control leg |
| **Activity** | | |
| *Moving* | V1 | Moving actively around |
| *Cowering* | V2 | Laying on the back or side and not moving (shock or dead) |
| *Standing* | V3 | Standing straight/still |
| **Interaction** | | |
| *Marker* | X1 | Interacting with red wire loop |

*Results*

Injured ants engaged in allogrooming more frequently and for longer durations ($\chi^2 = 15.17$; $p = 0.0001$; $\chi^2 = 22.42$; $p < 0.0001$, respectively) and showed a higher occurrence of cowering ($\chi^2 = 7.87$; $p = 0.005$) than control ants. Conversely, injured ants spent less time selfgrooming ($\chi^2 = 4.06$; $p = 0.05$) and engaged in selfgrooming less frequently ($\chi^2 = 4.98$; $p = 0.03$). They were also less frequently moving ($\chi^2 = 13.13$; $p = 0.0003$) or standing ($\chi^2 = 5.6$; $p = 0.02$) compared to control ants (Fig. S13). The duration of allogrooming was linked to an interaction between treatment and climate ($\chi^2 = 7.80$; $p = 0.005$; Fig. S14). Across climatic gradients, allogrooming duration ($\chi^2 = 15.28$; $p = 0.0001$) and allogrooming occurrence ($\chi^2 = 7.36$; $p = 0.007$) were negatively associated with PC1 climate. In contrast, both moving occurrence ($\chi^2 = 3.84$; $p = 0.05$) and standing occurrence ($\chi^2 = 3.88$; $p = 0.05$) increased with PC1 climate. Higher parasite prevalence was associated with reduced allogrooming duration ($\chi^2 = 12.75$; $p = 0.0004$), while the occurrence of standing increased with parasite prevalence ($\chi^2 = 5.32$; $p = 0.02$).

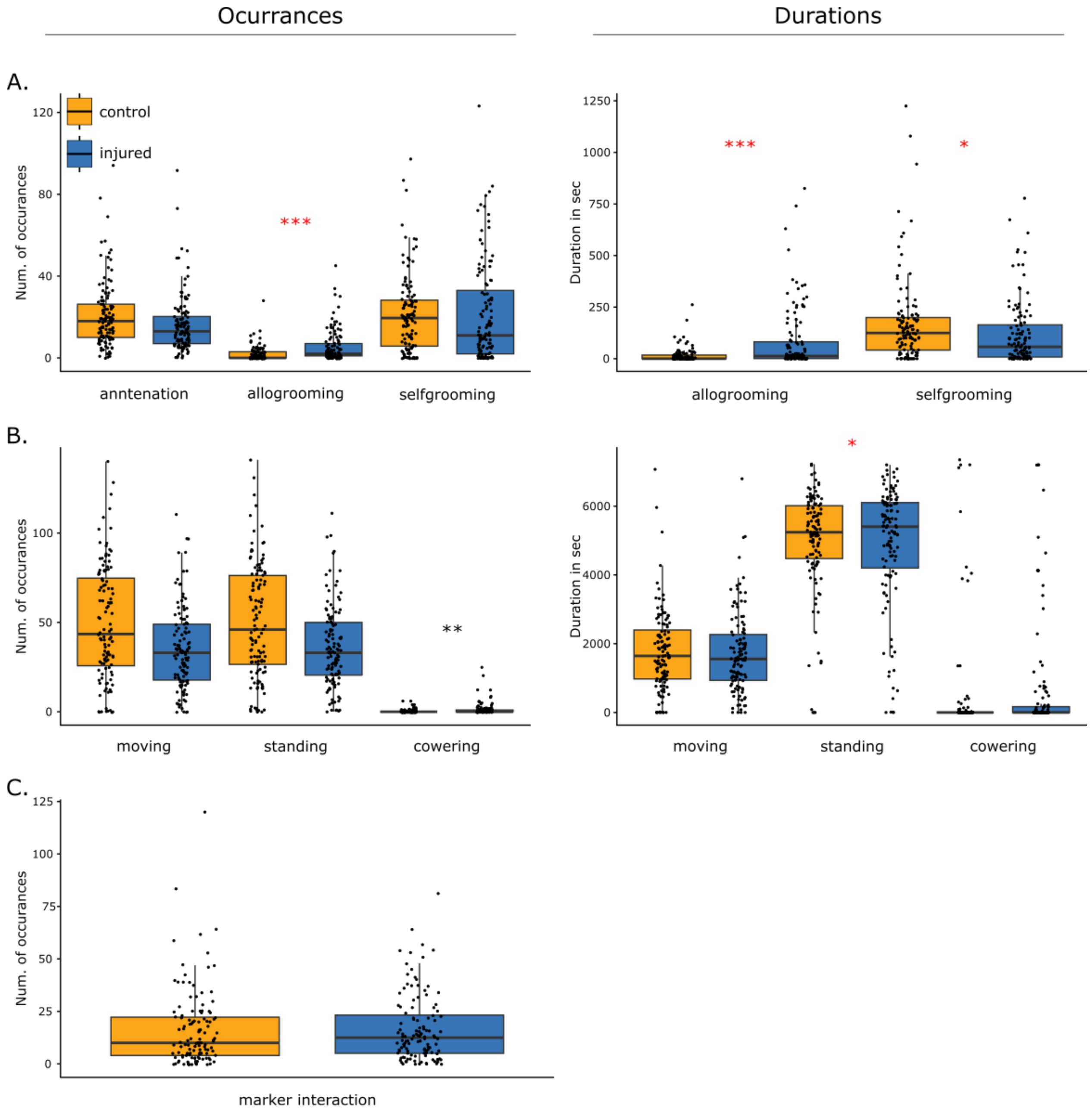


**Figure S13. Behavioural responses to injury.** Boxplots of occurrences and durations of (A) social and grooming behaviour, (B) active behaviour, and (C) interaction with the wire loop. Significant differences ($p < 0.05$) between control and experimental ants are depicted via red asterisks.

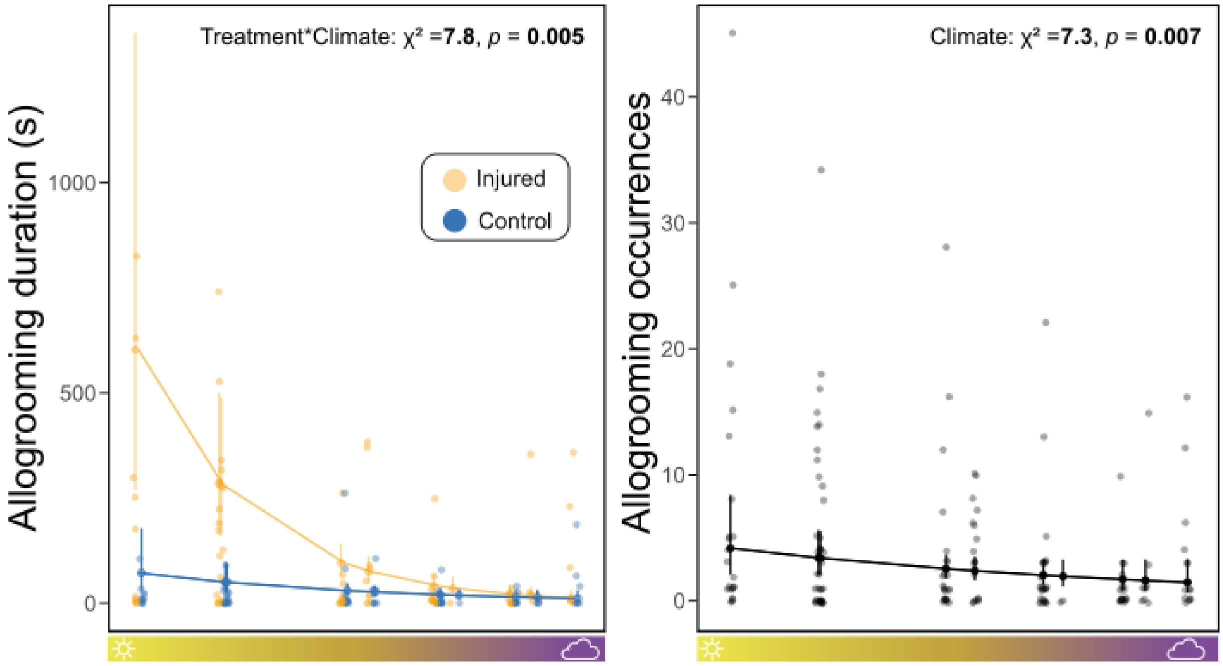


**Figure S14. Effects of climate on allogrooming duration and occurrence.** Allogrooming duration was linked to an interaction between treatment and climate ($\chi^2 = 7.80$; $p = 0.005$). Across climatic gradients, allogrooming duration ($\chi^2 = 15.28$; $p = 0.0001$) and allogrooming occurrence ($\chi^2 = 7.36$; $p = 0.007$) were negatively associated with PC1 climate.

*Discussion and Conclusion*

This study examined social and grooming behaviours in response to ant injury, considering the effects of climate and parasite prevalence while controlling for colony size. Results showed that ants from warmer climates engaged in more frequent and prolonged wound grooming of injured nestmates, suggesting a positive effect of temperature on altruistic care. Notably, the duration of allogrooming was higher in injured ants compared to controls only in warmer regions. The presence of parasites in these regions, combined with the heightened risk of infection from higher average temperatures that lead to the proliferation of pathogens (Linder et al., 2008), may drive this increased care to minimise worker loss and also prevent the spread of potential microbial infections following parasite raids. However, our analysis also showed that higher parasite prevalence correlated with less allogrooming, possibly indicating that in highly parasitised environments, colonies may prioritise resources differently, ‘replacing’ injured ants via brood care rather than investing in wound care, given the higher risk of fatality from infections.
